# Comparison of insect and human cytochrome b561 proteins: Insights into candidate ferric reductases in insects

**DOI:** 10.1101/2023.09.01.555970

**Authors:** Jessica D. Holst, Laura G. Murphy, Maureen J. Gorman, Emily J. Ragan

## Abstract

Cytochrome b561 (cytb561) proteins comprise a family of transmembrane oxidoreductases that transfer single electrons across a membrane. Most eukaryotic species, including insects, possess multiple cytb561 homologs. To learn more about this protein family in insects, we carried out a bioinformatics-based investigation of cytb561 family members from nine species representing eight insect orders. We performed a phylogenetic analysis to classify insect cytb561 orthologous groups. We then conducted sequence analyses and analyzed protein models to predict structural elements that may impact the biological functions and localization of these proteins, with a focus on possible ferric reductase activity. Our study revealed three orthologous groups, designated CG1275, Nemy, and CG8399, and a fourth group of less-conserved genes. We found that CG1275 and Nemy proteins are similar to a human ferric reductase, duodenal cytochrome b561 (Dcytb), and have many conserved amino acid residues that function in substrate binding in Dcytb. Notably, CG1275 and Nemy proteins contain a conserved histidine and other residues that play a role in ferric ion reduction by Dcytb. Nemy proteins were distinguished by a novel cysteine-rich cytoplasmic loop sequence. CG8399 orthologs are similar to a putative ferric reductase in humans, stromal cell-derived receptor 2. Like other members of the CYBDOM class of cytb561 proteins, these proteins contain reeler, DOMON, and cytb561 domains. *Drosophila melanogaster* CG8399 is the only insect cytb561 with known ferric reductase activity. Our investigation of the DOMON domain in CG8399 proteins revealed a probable heme-binding site and a possible site for ferric reduction. The fourth clade includes a group of proteins with a conserved “KXXXXKXH” non-cytoplasmic loop motif that may be a substrate binding site and is present in a potential ferric reductase, human tumor suppressor cytochrome b561. This study provides a foundation for future investigations of the biological functions of cytb561 genes in insects.

## Introduction

The cytochrome b561 (cytb561) proteins are a family of transmembrane oxidoreductases that function as electron transporters by accepting a single electron from a cytoplasmic reducing agent – generally assumed to be ascorbate – and transferring it across a membrane to reduce a non-cytoplasmic substrate [1,2]. Known non-cytoplasmic substrates (electron acceptors) include ferric (Fe^3+^) ions, which are reduced to ferrous (Fe^2+^) ions, and monodehydroascorbate (MDHA), which is reduced to ascorbate [2]. Animal and plant species possess multiple cytb561 homologs, some with ubiquitous expression and others with restricted expression patterns [1–3]. Depending on the particular protein, cytb561s can be located in the plasma membrane, the membranes of organelles in the secretory or endocytic pathways, or multiple membrane locations [4–7].

The biological functions of most cytb561s are uncertain, although some have established roles in iron reduction or ascorbate regeneration [1]. The well-characterized human duodenal cytochrome b561 (Dcytb) is a ferric reductase that facilitates iron uptake at the duodenal brush border, and it is also suspected to play other physiological roles [8–10]. The structurally well-characterized *Arabidopsis thaliana* (plant) cytochrome b561 B-paralog (AtCytb561-B) has in vitro ferric-chelate reductase activity, but its biological functions are unknown [11–13]. Other cytb561 family members with in vitro ferric reductase activity include mammalian stromal cell-derived receptor 2 (SDR2), tumor suppressor 101F6 (TScytb), chromaffin granule cytb561 (CGcytb), and lysosomal cytb561 (Lcytb), as well as *Drosophila melanogaster* (fly) CG8399, and *A. thaliana* AtCytb561-A [6,11,12,14,15]. The physiological significance of these proteins’ ferric reductase activity is not clear; for example, although CGcytb can reduce ferric-chelates in vitro, it is thought to function in chromaffin granules by reducing MDHA to produce ascorbate, which is then used by dopamine β-hydroxylase (DBH) in the synthesis of norepinephrine [16,17]. It seems likely that at least some cytb561s are multifunctional proteins, and that substrate specificity may be influenced by the substrate availability where a particular cytb561 is located.

To date, two cytb561 structures have been solved: human Dcytb and plant AtCytb561-B [13,18]. These structures confirmed many predicted cyb561 structural elements. Cytb561s have six transmembrane helices, including four consecutive helices that form a cytb561 core domain [2,13,18]. Inside the core domain are two b-type heme groups located near each side of the membrane, which are coordinated by four universally conserved histidine residues: the first- and third-conserved histidines (H1, H3) coordinate the non-cytoplasmic heme group, while the second- and fourth-conserved histidines (H2, H4) coordinate the cytoplasmic heme group [2,19].

Most cytb561s are single-domain proteins in which the core domain encompasses transmembrane helices two through five (TM2 through TM5) [2,13,18]. An exception is the CYBDOM proteins, which are multidomain cytb561s characterized by having at least one extracellular DOMON domain upstream of the cytb561 domain [1]. The CYBDOM cytb561 core domain encompasses TM1 through TM4, rather than TM2 through TM5 [1]. Single-domain cytb561s are oriented in a membrane with their N- and C-termini exposed to the cytoplasm [2]. In contrast, CYBDOMs have an extracellular-facing N-terminus [1], and, given that they have an even number of transmembrane regions, must also have an extracellular-facing C-terminus. Single domain cytb561s and CYBDOMs have three core domain loops that occur in the following order: cytoplasmic, non-cytoplasmic, cytoplasmic [1]. The function of the DOMON domain(s) in CYBDOMs is unknown, but some members of the DOMON domain superfamily bind sugar or heme, and at least one heme-binding DOMON domain is involved in interdomain electron transfer [20]. Some CYBDOMs also have an extracellular, N-terminal adhesion module, such as the reeler domain in SDR2 and *D. melanogaster* CG8399 [1]. The function of the reeler domain in CYBDOMs is not known.

Our study aims to better understand the cytb561 family in insects, and we are particularly interested in the possibility that insect cytb561 proteins play a role in cellular iron uptake. Iron homeostasis in insects is poorly understood, and mechanisms of cellular iron uptake are still unknown [21–23]. Most iron in biological systems is in the insoluble ferric form, but the known iron transporters in animals are specific for ferrous ions; thus, for iron to enter an animal cell through a metal transporter, it typically is first reduced to its soluble, ferrous state [24–26]. In mammals, the confirmed ferric reductases are Dcytb and various STEAP proteins [8,27–29]. Insects do not have STEAP homologs [21]; therefore, members of the cytb561 family are the most promising ferric reductase candidates in insects.

Surprisingly little is known about insect cytb561 proteins. Only one, *D. melanogaster* Nemy, has been studied in depth [4,30]. Nemy participates in a peptide amidation process in neuroendocrine neurons and influences olfactory learning and memory [4,30]. The *D. melanogaster* CYBDOM, CG8399, is reducible by ascorbate and has in vitro ferric reductase activity, but its biological function is unknown [6,31]. Nemy and *D. melanogaster* CG1275, an uncharacterized cytb561, have been described as possible ferric reductases [22,32].

The primary aims of this project were to 1) perform a phylogenetic analysis of cytb561 genes from nine diverse insect species to identify orthologous groups, 2) compare their amino acid sequences and predicted protein structures to those of better-studied human cytb561s to reveal sequence motifs and structural elements that may be involved in the localization and biological functions of these proteins, and 3) predict which of the orthologous groups may function as ferric reductases in insects. Of the four cytb561 groups identified, the CG1275 and Nemy groups are similar to human Dcytb; the CG8399 proteins are similar to human SDR2; and a subset of proteins in the fourth group is similar to human TScytb. We found that CG1275 and Nemy proteins have conserved amino acid residues and predicted protein structures that are consistent with ferric reductase activity. CG8399 proteins have highly-conserved amino acid residues in cytoplasmic regions and the DOMON domain that may be involved in substrate binding and ferric reductase activity. TScytb-like proteins have a “KXXXXKXH” motif that may be involved in non-cytoplasmic substrate binding and potential ferric reductase activity.

## Materials and methods

### Identification of insect cytb561 protein sequences

For this study, we selected *D. melanogaster* as our model insect species, along with eight additional insect species of interest: *Anopheles gambiae* (mosquito), *Acyrthosiphon pisum* (aphid), *Apis mellifera* (bee), *Ctenocephalides felis* (flea), *Papilio xuthus* (butterfly), *Pediculus humanus corporis* (louse), *Tribolium castaneum* (beetle), and *Zootermopsis nevadensis* (termite). These species were chosen for their phylogenetic diversity and their association with high-quality genomic data.

We identified *D. melanogaster* cytb561 protein sequences using the FlyBase protein domain search tool with the query “Cytochrome b561” [33]. Of the nine resulting hits, eight possessed the four conserved heme-coordinating histidine residues characteristic of cytb561 family members: CG1275, Nemy, CG8399, CG10165, CG13077, CG13078, CG10337, and CG3592. The remaining sequence, CG6479, lacked these conserved histidines and was eliminated from our dataset.

To identify cytb561 sequences in the insect species of interest, we performed species-specific BLAST-P searches using the National Center for Biotechnology Information (NCBI) non-redundant database [34]. Eight *D. melanogaster* cytb561 amino acid sequences were used as queries: NP_728727.1 (CG1275), NP_725208.1 (Nemy), NP_611079.2 (CG8399), NP_609982.1 (CG10165), NP_609990.1 (CG13077), NP_609989.1 (CG13078), NP_609986.1 (CG10337), and NP_570039.1 (CG3592). Additional insect cytb561 sequences were identified using the NCBI Conserved Domain Database (CDD) [35]. The NCBI CDD search engine was populated with a combination of “insect species” and either “cyt_b561” or “cytochrome_b_N,” and a search was executed.

Isoforms and identical hits were removed from the dataset, resulting in one unique sequence for each gene. Of the 46 non-*D. melanogaster* sequences discovered, 36 resulted from BLAST-P searches and ten resulted from NCBI CDD queries. A sequence from *T. castaneum* (XP_008195477.1) was edited using RNA-seq data to remove an apparently misplaced intron, resulting in the addition of 53 C-terminal amino acids and a complete cytb651 core domain. The NCBI accession number and source of each sequence is listed in S1 Table.

Three sequences with unique properties were observed: XP_314065.2 (*A. gambiae*), XP_001950579.2 (*A. pisum*), and XP_001949276.1 (*A. pisum*). These three sequences were used as separate queries to conduct BLAST-P searches against the NCBI non-redundant database for the class Insecta to determine whether other insect species have proteins with similar sequences.

### Defining homologous regions of protein sequences

Prior to performing phylogenetic and sequence analyses, we identified the homologous region of each insect cytb561 protein. We used the HHpred server (Max Planck Institute Bioinformatics Toolkit webserver), which is a tool for protein remote homology detection that generates a multiple sequence alignment, predicts secondary structure, then creates a profile hidden Markov model that is compared with a target database [36]. We evaluated each insect and outgroup sequence using the HHpred PDB_mmCIF70_3_Mar server set to default parameters, including local alignment. This server references the Protein Data Bank (PDB) of known protein structures for possible templates, then generates results based on the probability percentage that the query shares a homologous region with the template. An estimated probability percentage of > 95% indicates that homology is nearly certain [36]. All insect and outgroup sequences were determined to share the highest probability percentage of homology (> 99%) with regions of Dcytb (CYBRD1, NP_079119.3, PDB 5ZLG). The predicted homologous region was not identical for all insect sequences; therefore, we identified the region that all insect sequences have in common and defined this as the common homologous region, which corresponds to Dcytb A44-T179. We used common homologous sequences for all analyses unless otherwise described. The common homologous region of each sequence is listed in S1 Table.

To compare insect cytb561s with the six known human cytb561 proteins, we used HHpred and the same methodology described above to define the common homologous regions of the following human sequences: NP_079119.3 (Dcytb; CYBRD1), NP_705839.3 (Lcytb; CYB561A3), NP_001017916.1 (CGcytb; CYB561), NP_001278213.1 (TScytb; CYB561D2), NP_872386.1 (CYB561D1), and NP_001347970.1 (SDR2; FRRS1).

HHPred analysis of the DOMON domains in insect CG8399 orthologs was performed. All insect DOMONs were predicted to have > 99% predicted homology with regions of the cytochrome domain in the *Phanerodontia chrysosporium* (fungus) cellobiose dehydrogenase (CDH) protein (PDB 1D7B) and the DOMON domain in the human dopamine beta-hydroxylase (DBH) protein (PDB 4ZEL).

### Multiple sequence alignments, sequence identities, and localization

Multiple sequence alignments were performed in Clustal Omega and visualized in JalView, where we applied a custom color scheme and obtained consensus sequence logos and conservation scores [37,38]. Accession numbers for insect sequences used in alignments are in S1 Table. Additional sequences used in alignments include Dcytb and *Arabidopsis thaliana* cytochrome B561-1 (AtCytb561-B, NP_198679.1, PDB 4O7G). To determine sequence identities, we used CLUSTAL Omega to generate multiple sequence alignments for each insect sequence with all *D. melanogaster* and human cytb561 sequences, then determined pairwise sequence identities through utilization of the M-View multiple alignment viewer [39].

Full-length insect cytb561 protein sequences were submitted to the SignalP-5.0 webserver for detection of predicted signal peptides, and they were analyzed by the DEEPLOC-1.0 webserver for prediction of subcellular protein localization [40,41]. Sequences were also investigated by eye to identify putative late-endosomal/lysosomal targeting signals in the C-terminal portion of the protein [42].

### Phylogenetic analysis

For our phylogenetic analysis, we selected an outgroup species, *Trichoplax adhaerens*, based on two qualifying criteria: *T. adhaerens* belongs to an ancient animal lineage near the base of the metazoan tree, and its cytb561 genes are homologous to insect cytb561 genes [43]. To identify an outgroup sequence, we evaluated the cytb561 genes from *T. adhaerens* and selected the sequence with the lowest identity to the *D. melanogaster* cytb561 sequences. The phylogenetic analysis was conducted using the MEGA version X software [44]. First, all common homologous insect cytb561 sequences and the outgroup cytb561 sequence were aligned in CLUSTAL Omega using the “Pearson/FASTA” output parameter; the resulting alignment was uploaded to MEGA X, and a rooted phylogenetic tree was created using the maximum likelihood (ML) method. To determine the optimal model for ML tree construction, we ran the “Find Best DNA/Protein Models (ML)” tool in MEGA X, which resulted in recommended parameters: the LG with freqs. (+F) substitution model, gamma distributed with invariant sites (G+I) rates among sites, and five discrete gamma [45]. We then used the additional remaining parameters: use all sites in cases of gaps and missing data, the Nearest-Neighbor-Interchange ML heuristic method of tree inference, the default NJ/BioNJ initial tree for ML, no branch swap filters, seven threads, and 1000 bootstrap replicates.

### AlphaFold structure analysis

AlphaFold Monomer v2.0 pipeline structure predictions for *D. melanogaster* CG1275 isoform A (AF-Q9I7U1-F1-model_v4.pdb), *D. melanogaster* Nemy isoform A (AF-Q7JR72-F1-model_v4.pdb), *D. melanogaster* CG8399 isoform A (AF-Q8MSU3-F1-model_v4.pdb), *D. melanogster* CG13077 isoform B (AF-M9PG89-F1-model_v4.pdb), and *A. gambiae* XP_3200673.4 (AF-Q7PZD0-F1-model_v4) were downloaded from UniProt and visualized in ChimeraX [46–48]. We removed amino acids 1-30 from the CG8399 structure prediction as they were predicted to be a signal peptide by SignalP 6.0 [49]. AlphaFold structures for CG1275 and Nemy were individually overlayed with Dcytb (PDB 5ZLG) using the MatchMaker feature in ChimeraX with default settings. The number of α-carbon pairs and their root-mean-square deviation (RMSD) was generated by ChimeraX. CG1275 (amino acids Pro100-Ile326) and Nemy (amino acids Cys52-Ser281) were displayed together with Dcytb (amino acids Tyr6-Pro230) to generate figures. Individual AlphaFold models were associated with multiple sequence alignments of same group proteins and colored in ChimeraX using AL2CO and a color gradient from cyan (low conservation score) to magenta (high conservation score) [50]. The heme- and ascorbate-binding residues were selected to calculate possible intermolecular hydrogen bonds using the H-bonds tool and to predict possible direct residue interactions with the Contacts tool in ChimeraX using default settings for overlap cutoff and H-bond allowance parameters. Potential hydrogen bonds were displayed in blue. Coulombic surface coloring was used to visualize surface charges in the potential ion-binding pocket of CG8399 [46].

### Analysis of DOMON and TScytb sequence motifs

Additional sequence analyses were conducted to investigate DOMON domain sequences and a “KXXXXKXH” motif in cytb561 proteins from diverse animal species. Tree of Life was referenced to choose a set of species for these analyses: *Homo sapiens* (human), *Mus musculus* (mouse), *Passer montanus* (sparrow), *Xenopus laevis* (frog), *Danio rerio* (zebrafish), *Octopus sinensis* (octopus), *Actinia tenebrosa* (sea anemone), *Helobdella robusta* (leech), and *Acanthaster planci* (starfish) [51]. To identify DOMON-containing sequences, the *D. melanogaster* CG8399 sequence (NP_611079.2) was used as a BLAST-P query to search NCBI non-redundant databases; to identify “KXXXXKXH”-containing sequences, the query was *A. gambiae* sequence XP_320673.4. CLUSTAL Omega was then used to generate sequence alignments that allowed us to evaluate amino acid conservation of the DOMON domains and “KXXXXKXH” motifs and to determine the placement of the “KXXXXKXH” motif within the protein [39].

## Results and Discussion

### Phylogenetic analysis of insect cytb561 proteins

#### Insects have four distinct cytb561 groups

While a limited number of insect cytb561 sequences have been included in broader phylogenetic studies [3,19], we were interested in classifying the cytb561 family across diverse insect species. We identified 54 genes that encode cytb561 family members from nine insect species in eight orders: Blattodea (*Z. nevadensis*), Coleoptera (*T. castaneum*), Diptera (*A. gambiae* and *D. melanogaster*), Hemiptera (*A. pisum*), Hymenoptera (*A. mellifera*), Lepidoptera (*P. xuthus*), Psocodea (*P. humanus corporis*), and Siphonaptera (*C. felis*) (S1 Table). All investigated species have between three *(P. humanus corporis*) and nine (*P. xuthus*) cytb561 genes (S1 Table).

Using the HHpred PDB server for protein remote homology detection, we defined a homologous region (core domain) shared by all of the insect cytb561s [36] (S1 Table). The ∼130 amino acid long core domain sequences were then used to generate a multiple sequence alignment (S1 Fig). From this multiple sequence alignment, we generated a phylogenetic tree and found strong support for four cytb561 groups: three orthologous groups and a fourth group that lacks one-to-one orthologs (Fig 1). For simplicity, we will refer to all insect sequences that cluster in the three orthologous groups according to the representative *D. melanogaster* protein name: CG1275, Nemy, or CG8399. The remaining 25 sequences will be referred to as Group 4.

**Fig 1.**
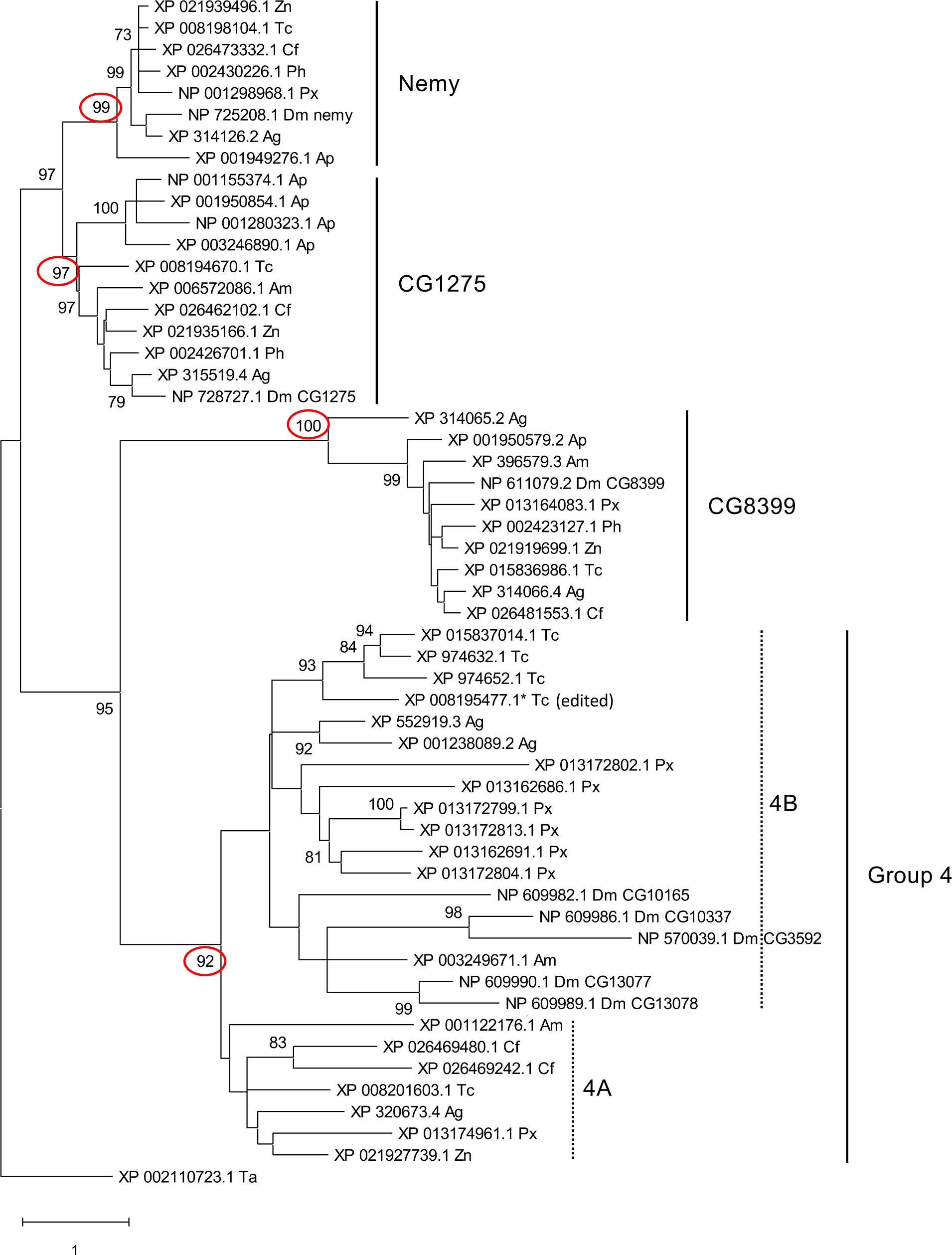
Phylogenetic tree of insect cytb561 proteins. A phylogenetic analysis of cytb561s from nine species of insects (*A. gambiae [Ag]*, *A. mellifera [Am]*, *A. pisum [Ap]*, *C. felis [Cf]*, *D. melanogaster [Dm]*, *T. castaneum [Tc]*, *P. humanus corporis [Ph]*, *P. xuthus [Px]*, and *Z. nevadensis [Zn])* was performed with the maximum likelihood method. An outgroup sequence from *T. adhaerens (Ta)* was used for rooting the tree. Bootstrap values greater than 70% are shown. Three orthologous groups (with one-to-one orthologs in most or all species) were identified: CG1275, Nemy, and CG8399. The remaining genes formed a fourth group that has no one-to-one orthologs but includes many species-specific gene expansions. The bootstrap values associated with the four clades are marked with red circles. A branch length scale bar indicates one substitution per site. Of the Group 4 sequences, seven are more similar to human TScytb than any *D. melanogaster* cytb561s and are referred to as subgroup 4A, whereas the remaining Group 4 proteins belong to subgroup 4B. (Note that the 4A and 4B subgroups are based on sequence comparisons rather than phylogenetic relationship.)

#### CG1275 and Nemy

The CG1275 and Nemy groups are closely related, as evidenced by bootstrap support of a CG1275-Nemy cluster and the short branch lengths for the CG1275 and Nemy groups (Fig 1). CG1275 proteins were identified for eight of the nine insect species (Fig 1; S1 Table). We did not identify a CG1275 ortholog in the lepidopteran insect *P. xuthus*, and we were unable to find a CG1275 sequence in any other species from the order Lepidoptera (butterflies and moths). Of the species analyzed, *A. pisum* was the only insect with a CG1275 gene expansion (Fig 1; S1 Table). We identified Nemy proteins in eight of the species (Fig 1; S1 Table), but we did not find a Nemy sequence from *A. mellifera* or any other species in the order Hymenoptera (bees, wasps, ants, and similar insects). A previously published phylogenetic analysis of eukaryotic cytb561s suggested that CG1275 and Nemy from *D. melanogaster* and *A. gambiae* may be orthologous to human Dcytb, CGcytb, and Lcytb, but a lack of statistical support for the clade makes the phylogenetic relationship of CG1275 and Nemy with Dcytb, CGcytb, and Lcytb uncertain [19].

#### CG8399

CG8399 is the only orthologous group with sequences from all nine insect species. This cluster includes ten proteins, nine of which contain multiple domains – reeler, DOMON(s), and cytb561 – consistent with some members of the CYBDOM protein family (Fig 1; S1 Table) [1]. *A. gambiae* was the only species with a second CG8399-like protein (XP_314065.2); this sequence differs from other CG8399s in that it is a single-domain cytb561 lacking both reeler and DOMON domains. A previously published phylogenetic analysis of eukaryotic cytb561s demonstrated that *D. melanogaster* and *A. gambiae* CG8399 proteins are orthologous to SDR2 in humans, other animals, and plants [19].

#### Group 4

The remaining 25 insect genes populate a large clade (Group 4) with a bootstrap value of 92 (Fig 1). Group 4 has multiple species-specific clusters with significant bootstrap values, suggesting that Group 4 genes were more likely than other insect cytb561 genes to undergo species-specific expansions (Fig 1). We did not identify any Group 4 genes in *P. humanus corporis* or *A. pisum* (Fig 1).

### Sequence identity between human and insect cytb561s

Human cytb561s have been more widely studied than insect cytb561s, and human Dcytb is one of only two cytb561 structures that have been solved [18]. We were interested in determining the sequence similarities between the 54 insect cyb561s and the six human cytb561s (Dcytb, CGcytb, Lcytb, SDR2, TScytb, and CYB561D1) to learn which of the better-characterized human cytb561s is most similar toeach of the insect cytb561s. For this analysis, we identified the common homologous region of each insect and human cytb561 sequence, then determined their pairwise identities. CG1275 and Nemy proteins had highest sequence identity with Dcytb, CGcytb, or Lcytb (32.4%-49.3%), and CG8399s had highest sequence identity with SDR2 (36.5%-43.4%) (Table 1). These results suggest that CG1275 and Nemy proteins may have physiological similarities with Dcytb, CGcytb, and/or Lcytb, and are consistent with the finding that CG8399s are orthologous to SDR2 [19]. Of the 25 Group 4 cytb561s, 23 had highest sequence identity with TScytb (21; 18.8%-33.1%) or CYB561D1 (2; 22.0%-25.7%) (Table 1). Seven Group 4 proteins are distinct from all other insect cytb561s in that they had higher sequence identity with a human cytb561 (TScytb) than with any of the *D. melanogaster* cytb561s; we will refer to this subset of proteins as Group 4A and the remaining Group 4 proteins as Group 4B (Table 1; Fig 1; S2 Table). A previous phylogenetic study found that the *A. gambiae* gene in Group 4A clustered with human and other TScytbs rather than other insect-specific sequences, but there was a lack of bootstrap support for this cluster; thus, the phylogenetic relationship between Group 4A genes and TScytb is unclear [19].

**Table 1.**
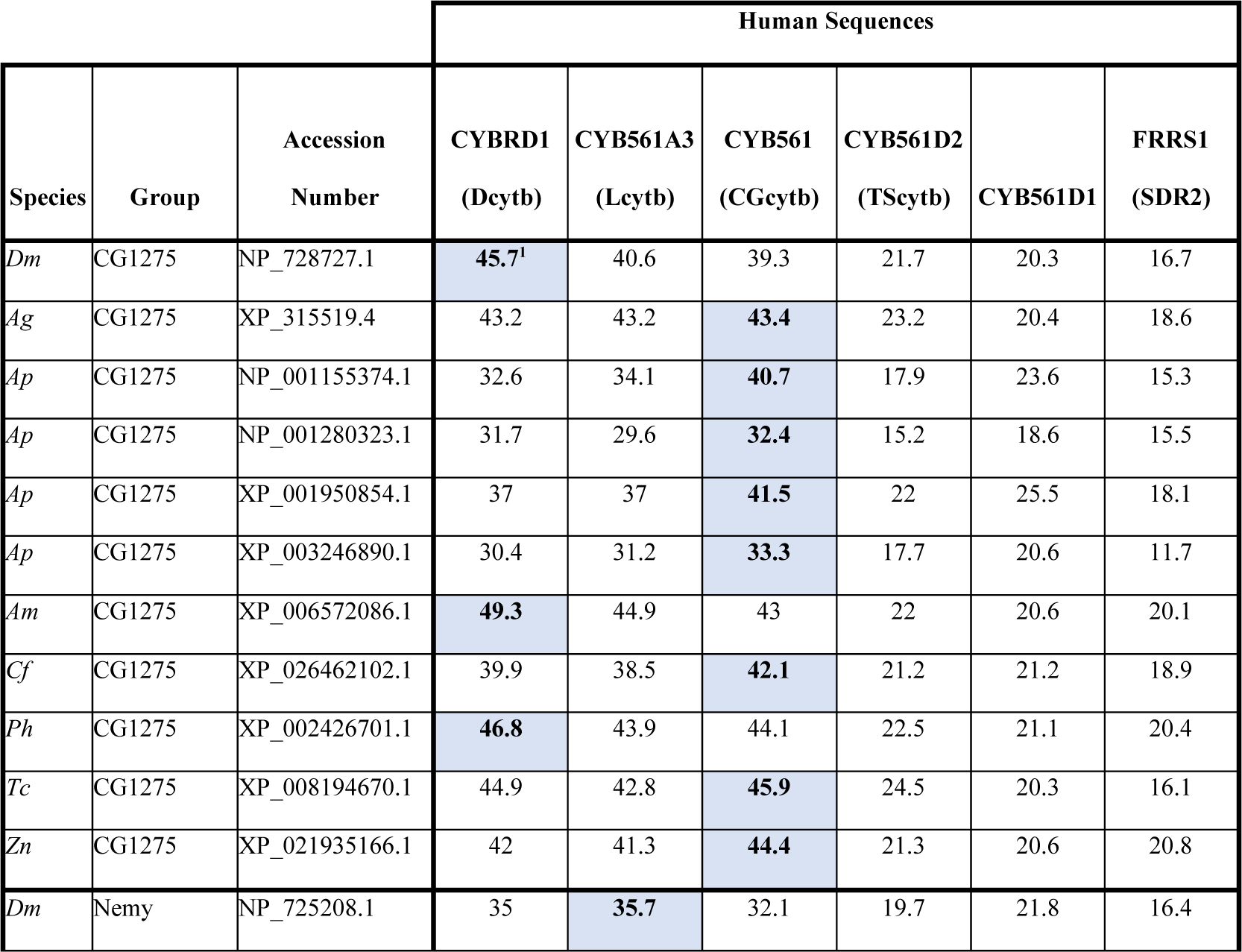
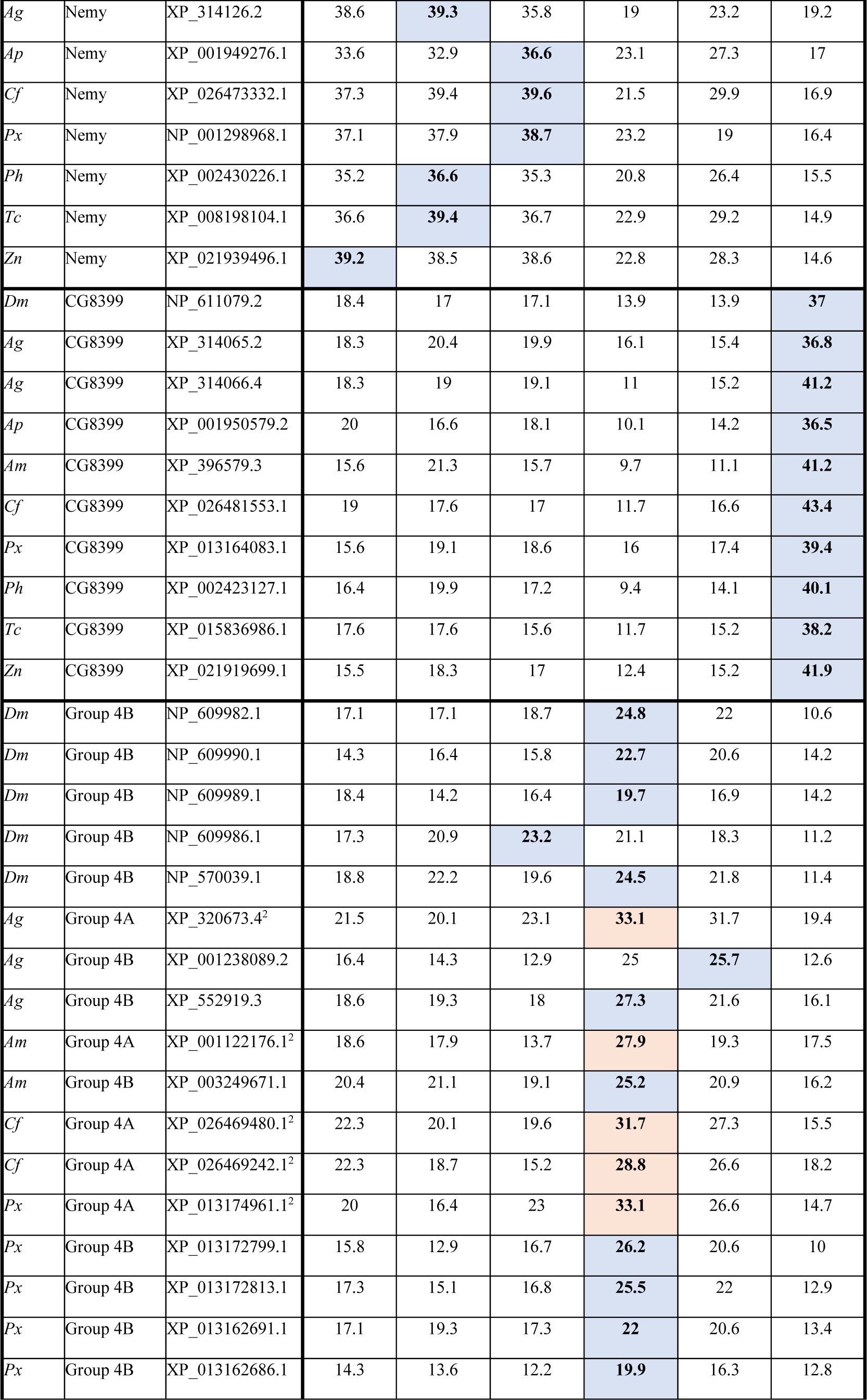
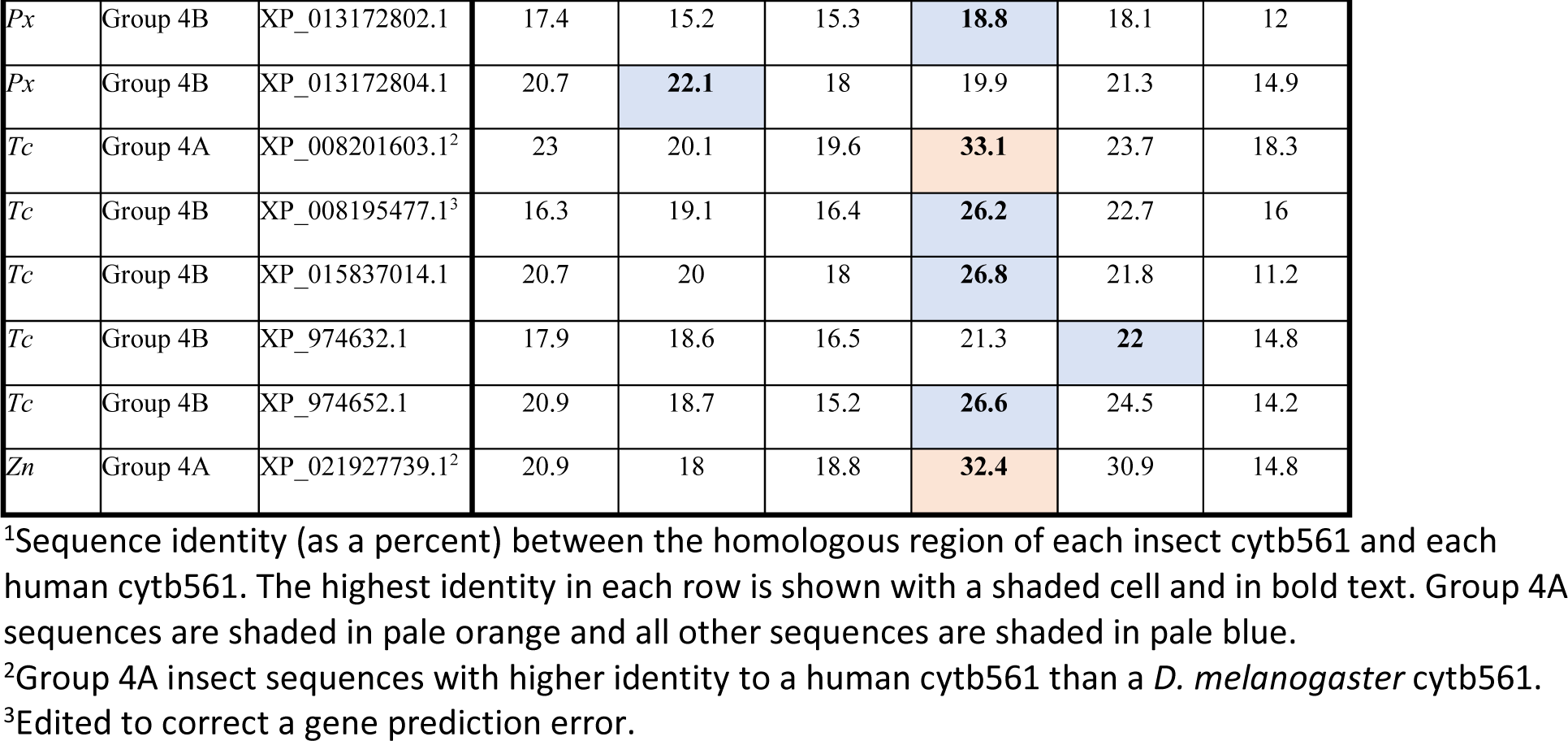
Sequence identity between insect and human cytb561s.

### Analysis of insect cytb561 domains

#### Locations of conserved amino acid residues

We used amino acid sequence alignments to identify conserved amino acid residues in insect cytb561 domains. To facilitate comparisons of residues from diverse sequences, we denote residues as the number of positions they are upstream (−) or downstream (+) of one of the four conserved heme-coordinating histidine residues; for example, the residue immediately upstream of H1 is referred to as H1-1. To identify amino acid residues that are conserved in all four insect cytb561 groups, we aligned the ∼130 amino acid core domain sequences of all 54 insect species and found that only the heme-coordinating histidines, H1-H4, were completely conserved (S1 Fig). Note that for the CG1275, Nemy, and Group 4 proteins, H1-H4 are in TM2-TM5, whereas in the CG8399 proteins, H1-H4 are in TM1-TM4. In addition to the four positions containing conserved histidines, 11 positions have highly-conserved residues (with Jalview conservation scores of 8 or 9), including seven with small residues (Gly, Ala, and Ser), and four with larger hydrophobic residues (S1 Fig). Five of the seven small residue positions were previously noted [3], including two that are present in 106 eukaryotic cytb561s (H1+7, H4+4) [19]. To identify amino acid conservation within specific insect cytb561 groups, we separated the core domain alignment into five group-specific alignments and then used Jalview to generate group-specific consensus logos (Fig 2) [38]. These consensus logos reveal patterns of conservation within groups and allow residues at specific locations to be compared across groups.

**Fig 2.**
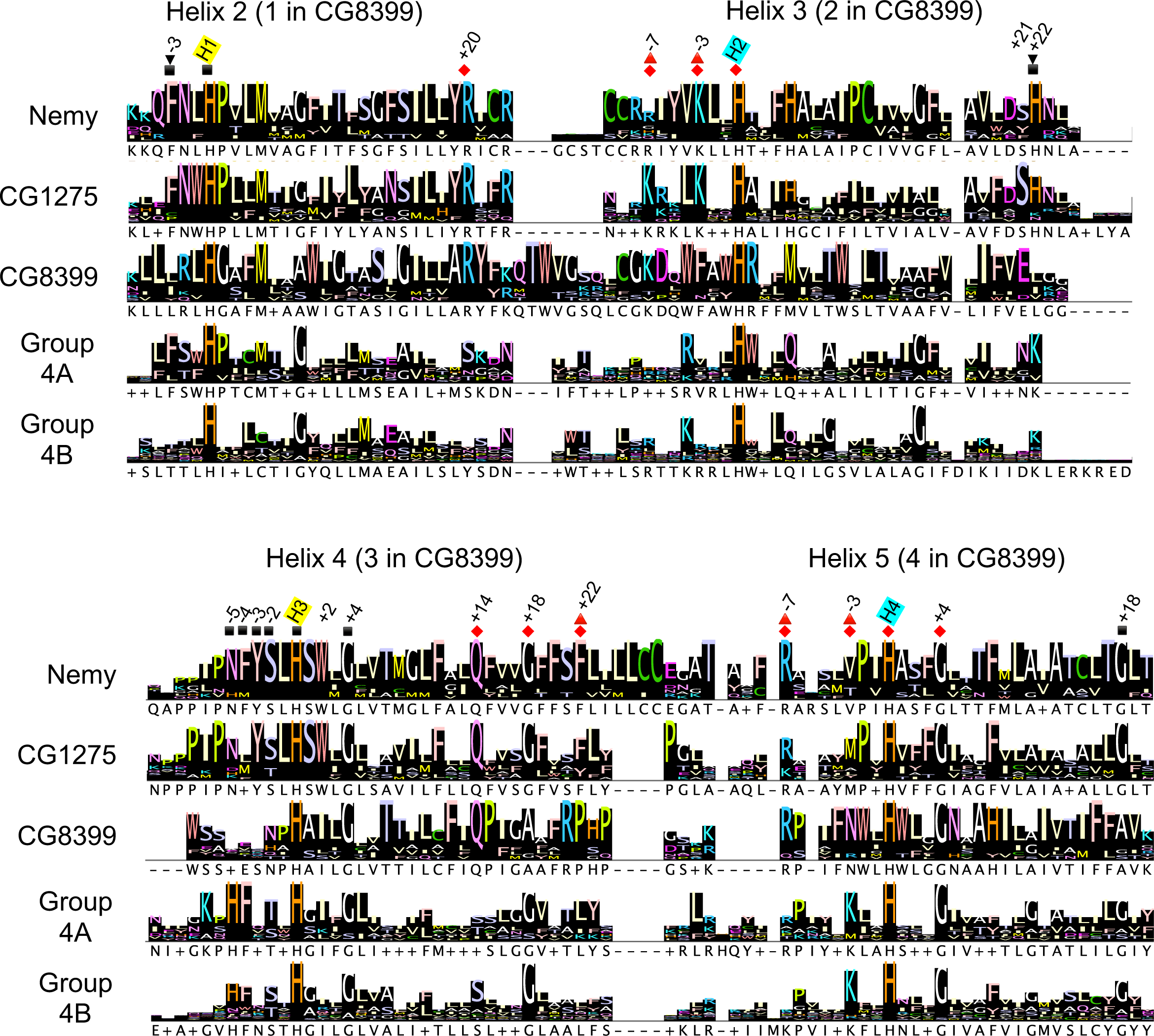
Group consensus logos for the cytb561 core domain. Group-specific alignments were used to create group consensus sequences in Jalview. The four conserved histidines are labeled H1-H4 with yellow highlighting for the non-cytoplasmic histidines and blue highlighting for the cytoplasmic histidines. The positions of amino acid residues of interest are labeled relative to the nearest conserved histidine. If the corresponding residue in Dcytb was predicted to contact a heme or ascorbate, that position is indicated by a symbol: non-cytoplasmic heme (square), non-cytoplasmic ascorbate (upside-down triangle), cytoplasmic heme (red diamond), or cytoplasmic ascorbate (red triangle). The Jalview consensus rows show the most common residue at that position listed below the logo; a + is used where there are two or more residues with the highest percentage and a – is used to show gaps.

To analyze the conserved amino acids in the context of the protein structure, we used ChimeraX to overlay a protein alignment of the CG1275, Nemy or CG8399 groups with its representative *D. melanogaster* AlphaFold protein model (Fig 3). For Group 4B proteins, we selected the *D. melanogaster* CG13077 AlphaFold model (Fig 3). Since *D. melanogaster* does not have a Group 4A protein, we used the *A. gambiae* Group 4A (XP_320673.4) AlphaFold model (Fig 3). The degree of conservation is shown by gradient coloring. By analyzing conservation information from the structural models as well as the group-specific consensus logos, we determined that the insect cytb561 proteins, with the exception of the Group 4B proteins, have relatively high group sequence conservation in the regions near non-cytoplasmic heme-coordinating H1 and H3 (Figs 2 and 3). Additionally, CG8399 proteins have relatively high conservation in the regions preceding the cytoplasmic heme-coordinating H2 and H4 (Figs 2 and 3).

**Fig 3.**
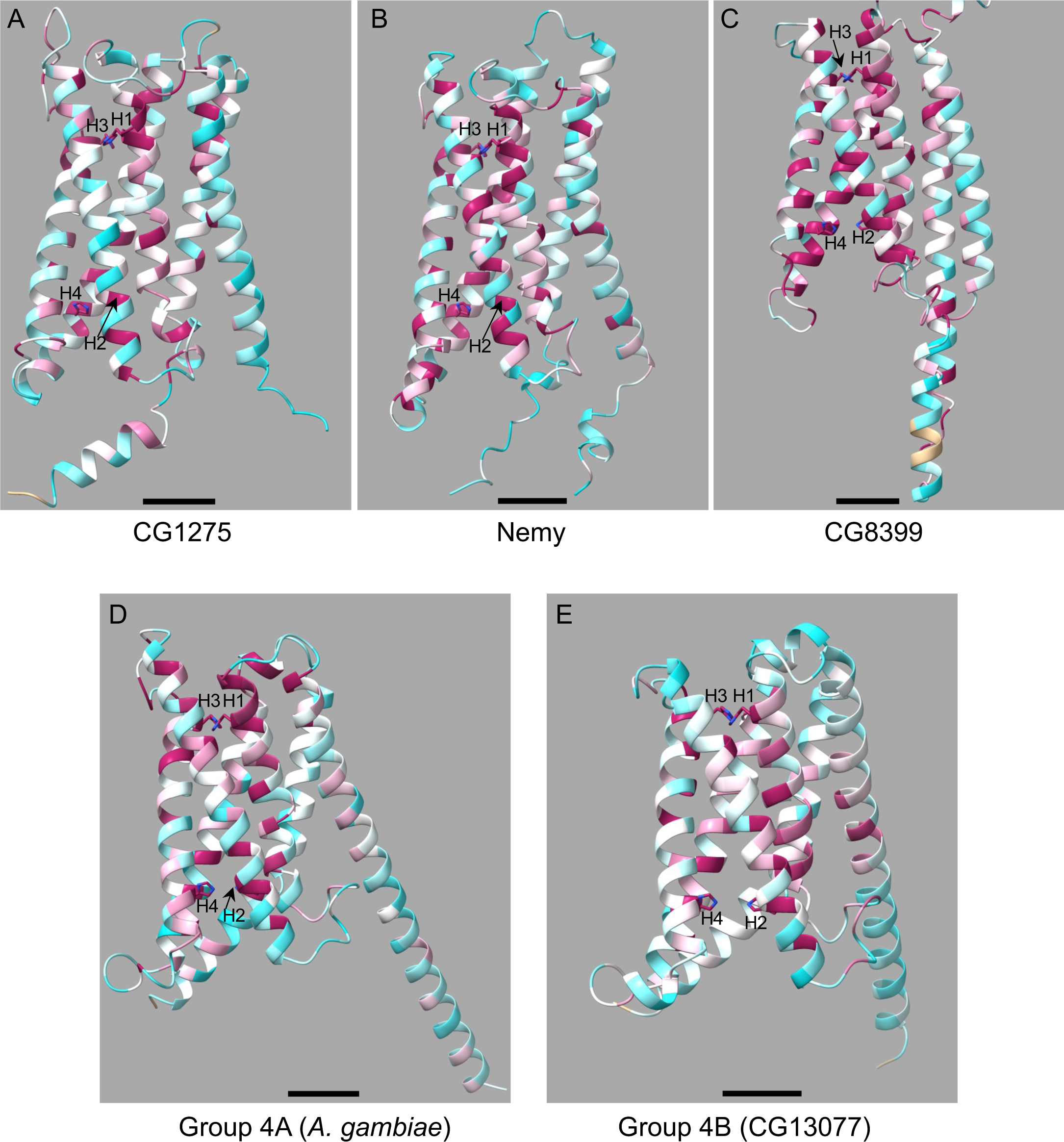
Insect cytb561 models with sequence alignment overlay demonstrate residue conservation. AlphaFold models of mature proteins are shown. Sequence conservation was determined by ChimeraX using AL2CO conservation scores. The model is gradient colored with high conservation (+1.5 score) in magenta and low conservation (−1.5 score) in cyan; sequence gaps are shown in tan. Conserved histidines H1-H4 (not all visible in all panels) are shown in stick form with nitrogen in blue. The models are oriented with the cytoplasmic side of the protein at the bottom. Note that the amino- and carboxyl-termini for all models except the CG8399 model are near the bottom of the panels. Locations of H1-4 are indicated. Scale bars depict 10 Å. A) *D. melanogaster* CG1275 (Lys95-Glu340) aligned with the CG1275 sequences. B) *D. melanogaster* Nemy (Lys40-Ile290) aligned with the Nemy sequences. C) *D. melanogaster* CG8399 (Ser402-Gly645) aligned with the multi-domain CG8399 sequences. The reeler and DOMON domains are omitted. D) *A. gambiae* XP_320673.4 (all residues) aligned with the Group 4A sequences. E) *D. melanogaster* CG13077 (all residues) aligned with the Group 4B sequences.

### General comparisons of insect cytb561 proteins with Dcytb and AtCytb561-B

This is the first large scale analysis of cytb561 proteins since X-ray crystallography studies identified heme- and ascorbate-binding residues in human Dcytb and *A. thaliana* AtCytb561-B, and a metal ion-coordinating residue in Dcytb [13,18]. We wanted to leverage this structural information to predict whether conserved residues in insect cytb561s might participate in heme, ascorbate, and/or iron-binding. Amino acid sequence alignments and structural models were used to compare insect cytb561 proteins with Dcytb and Atcyb-B. Initial comparisons were restricted to CG1275 and Nemy proteins because of their high similarity to Dcytb and AtCytb561-B; once key residues were identified in CG1275 and Nemy, we used sequence alignments to identify the corresponding residues in the CG8399 and Group 4 proteins. References to specific residue numbers are based on Dcytb unless otherwise noted.

We first created an alignment of Dcytb, AtCytb561-B, CG1275, and Nemy sequences (Fig 4). From this alignment, which extends beyond the core domain, we identified 23 completely conserved residues and an additional 23 residues that are highly conserved across Dcytb, AtCytb561-B, CG1275, and Nemy sequences (Fig 4). To identify any previously unspecified Dcytb residues that are likely to be important for heme or ascorbate binding, we used ChimeraX to determine predicted protein contacts with each heme and ascorbate group in the Dcytb ascorbate-bound structure (PDB 5ZLG) (Fig 4; S3 Table). (Note that Dcytb was selected for this analysis rather than AtCytb561-B for multiple reasons: insects are more closely related to humans than plants, Dcytb is a ferric reductase, and the Dcytb crystal structure contains a metal ion.)

**Fig 4.**
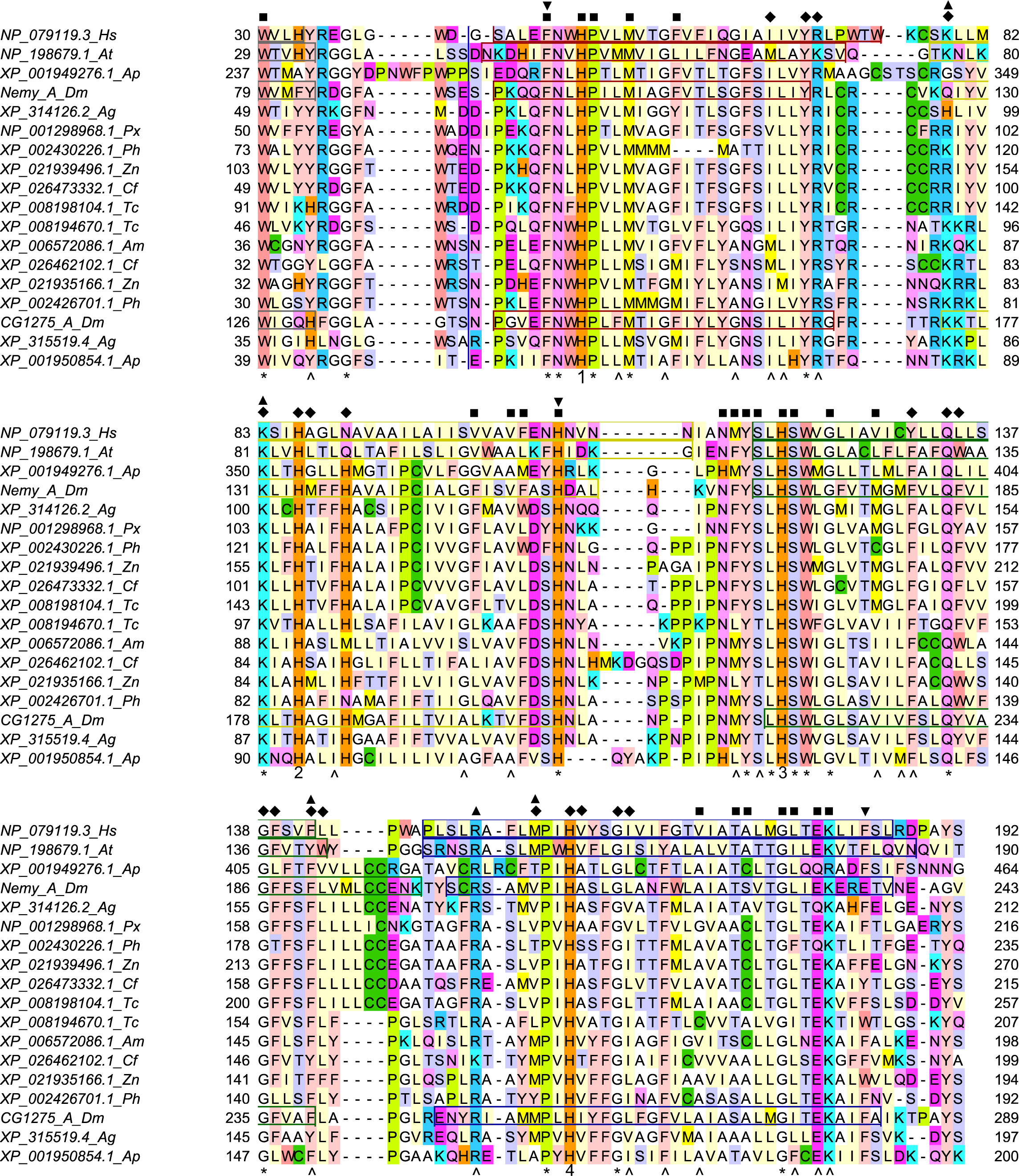
Alignment of Dcytb, AtCytb561-B, Nemy, and CG1275 protein sequences. The two cytb561 sequences with known structure, human Dcytb (PDB 5ZLG) and *A. thaliana* AtCytb561-B (PDB 4O7G), were aligned with CG1275 and Nemy sequences from *A. gambiae (Ag)*, *A. mellifera (Am)*, *A. pisum (Ap)*, *C. felis (Cf)*, *D. melanogaster (Dm)*, *T. castaneum (Tc)*, *P. humanus corporis (Ph)*, *P. xuthus (Px)*, and *Z. nevadensis (Zn)*. (Note that only the *A. pisum* CG1275 sequence most similar to Dcytb was included.) Alpha helices from crystal structures (Dcytb, AtCytb561-B) or predicted in AlphaFold models (Nemy, CG1275) are shown in boxes: helix 1 carboxyl end (gray), helix 2 (red), helix 3 (yellow), helix 4 (green), and helix 5 (blue). Conserved residues are noted under the alignment (* indicates completely conserved, the four strictly conserved histidines are labeled with their number rather than *, and ^ indicates highly conserved with a Jalview conservation score of 8 or 9). Predicted Dcytb heme-contacting residues are labeled with a square (non-cytoplasmic heme) or diamond (cytoplasmic heme), and ascorbate-contacting residues are labeled with an upside-down triangle (non-cytoplasmic ascorbate) or triangle (cytoplasmic ascorbate). A blue vertical line indicates the position of 51 amino acids found only in *A. pisum* XP_001949276.1, which were omitted from the figure. Coloring by amino acid residue: basic residues in blue (Arg) or cyan (Lys); acidic residues (Asp and Glu) in dark pink; amide residues (Asn and Gln) in light pink; hydroxylic residues (Ser and Thr) in pale purple; aromatic residues in light peach (Phe and Tyr) or dark peach (Trp); sulfur-containing residues in dark green (Cys) or light green (Met); His in orange; Pro in yellow; Ile, Leu, and Val in pale yellow; and Ala and Gly in white.

Next, we compared structural models of *D. melanogaster* CG1275 and Nemy with the solved structure of Dcytb. To investigate, we overlayed AlphaFold models for *D. melanogaster* CG1275 and Nemy with the ascorbate-bound structure of human Dcytb. The AlphaFold models have very high confidence for the majority of residues in the six transmembrane helices and their connecting loops (S4 Table), and the alpha carbons of the two insect structures aligned well over most of the Dcytb sequence (Fig 5) [47,48,52]. Although the AlphaFold cytb561 models do not include the two bound heme groups and the two bound ascorbates, overlaying the AlphaFold structures with Dcytb showed how these molecules may fit within the CG1275 and Nemy structures (Fig 5) [18].

**Fig 5.**
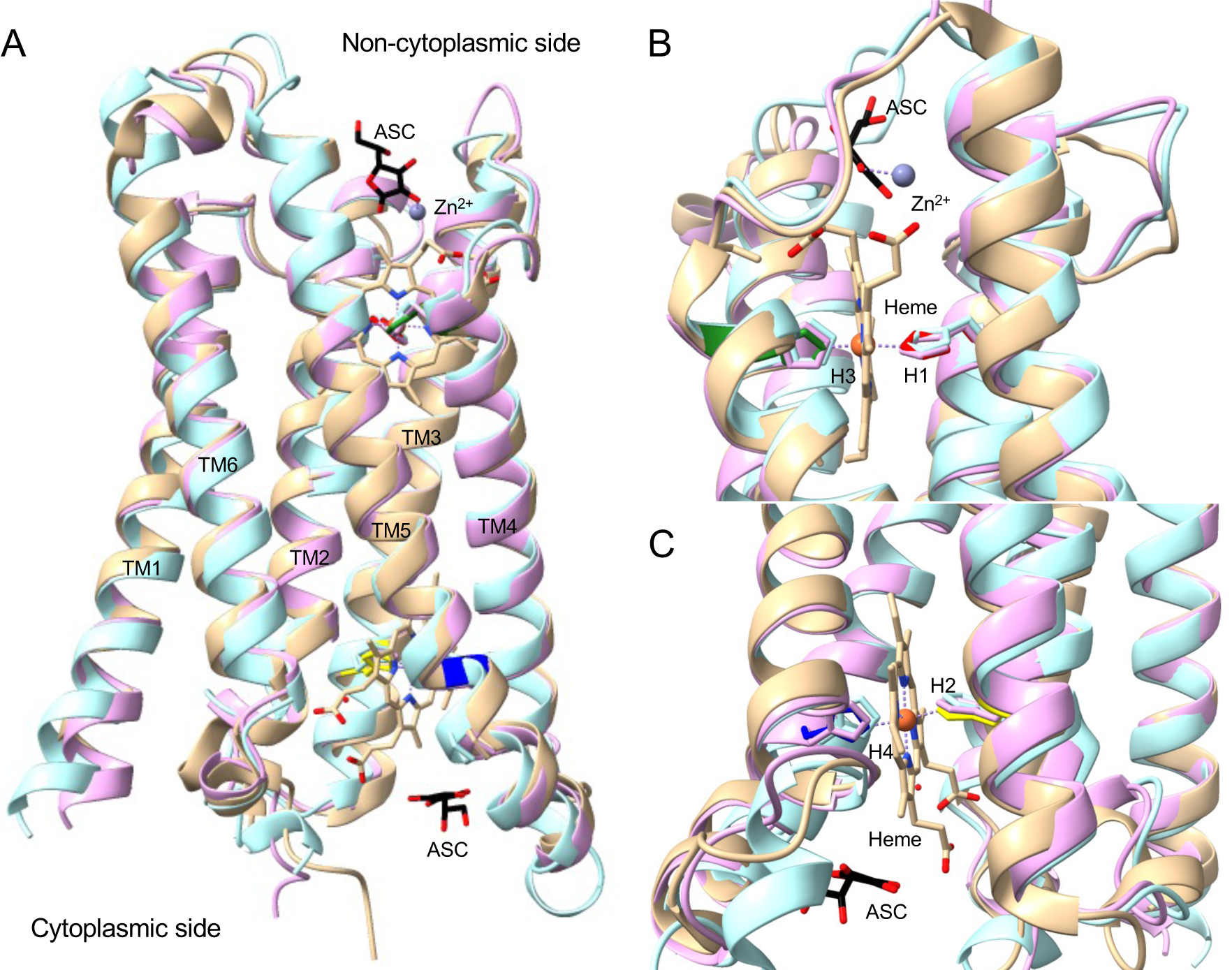
*D. melanogaster* CG1275 and Nemy models overlayed on Dcytb heme- and ascorbate-bound structure. AlphaFold structures of *D. melanogaster* CG1275 and Nemy were overlayed with Dcytb (PDB 5ZLG) using Chimera Matchmaker. Transmembrane helices are labeled (TM1-TM6), with Dcytb in tan, CG1275 in pale pink, and Nemy in pale blue. The four conserved histidines are shown in stick form; in the Dcytb structure, H1 is in red, H2 in yellow, H3 in green, and H4 in dark blue. Ascorbate (ASC) in black stick, heme in tan stick (with iron shown as orange spheres), and zinc (Zn^2+^) ion coordination in purple are from the 5ZLG structure. Stick side chains, ascorbate, and heme have oxygen in red and nitrogen in blue. The alpha carbon RMSD for CG1275 with 5ZLG is 0.679 Å for the 208 pruned atom pairs and 1.458 Å across all 222 pairs. The alpha carbon RMSD for Nemy with 5ZLG is 0.990 Å for the 188 pruned atom pairs and 2.657 Å across all 222 pairs. A) Overlay of *D. melanogaster* CG1275 and Nemy with Dcytb. B) Non-cytoplasmic ascorbate binding, zinc coordination, and heme coordination with H1 (red) and H3 (green) shown in stick form. C) Cytoplasmic ascorbate binding and heme coordination with H2 (yellow) and H4 (dark blue) shown in stick form.

### Conserved insect cytb561 residues likely to be involved in heme positioning

We were interested in exploring whether residues involved in heme placement in Dcytb and AtCytb561-B have similar roles in insect cytb561 proteins. Our Dcytb protein contact analysis identified 47 residues that make at least one predicted contact with a heme group (S3 Table), including 32 residues that are completely or well conserved in all CG1275 and Nemy sequences, as well as in AtCytb561-B (Fig 4). Many of these 32 residues are predicted to make contacts with a heme group in *D. melanogaster* CG1275 and Nemy protein models overlayed with Dcytb (S3 Table). In particular, the H1-H4 side chains are positioned very similarly to the heme-coordinating H1-H4 side chains in the Dcytb structure (Figs 5B and C). Amino acid residues that were predicted to interact with a heme group or participate in heme positioning, as well as their degree of conservation in the four insect cytb561 groups, are described below.

#### Cytoplasmic heme

The cytoplasmic heme group in Dcytb directly interacts with three amino acid residues: H2, H4, and Arg70 (H1+20) [18]. Arg70 forms a salt bridge with the heme A-propionate through its epsilon nitrogen [18]; a similar salt bridge is formed by a lysine at H1+20 in AtCytb561-B (Lys71) [13]. An arginine at H1+20 is completely conserved in all CG1275, Nemy, and CG8399 sequences (Fig 2) and likely interacts with the cytoplasmic heme A-propionate (Fig 6A). The residue at H1+20 tends to be a serine or threonine in Group 4A proteins (Fig 2), and 22 out of 25 Group 4B sequences also have a polar residue at H1+20 (S1 Fig). Serine is a much smaller residue than arginine, but in Dcytb the gamma carbon of Arg70 is only ∼3.2 Å from the heme A propionate; therefore, a serine or threonine residue could potentially form a hydrogen bond with a heme carboxylate group if properly positioned. Outside of the Dcytb core domain, at the end of helix 6, Lys225 forms two potential hydrogen bonds with the cytoplasmic heme D propionate (Fig 6A; S3 Table). This helix 6 lysine is conserved in CG1275 sequences and is a basic residue in 5 out of 7 Nemy sequences, but it is replaced with a proline in *D. melanogaster* and *A. gambiae* Nemy. (Comparisons of Lys225 with CG8399 and Group 4 sequences were not attempted because helix 6 is outside of their homologous regions.)

**Fig 6.**
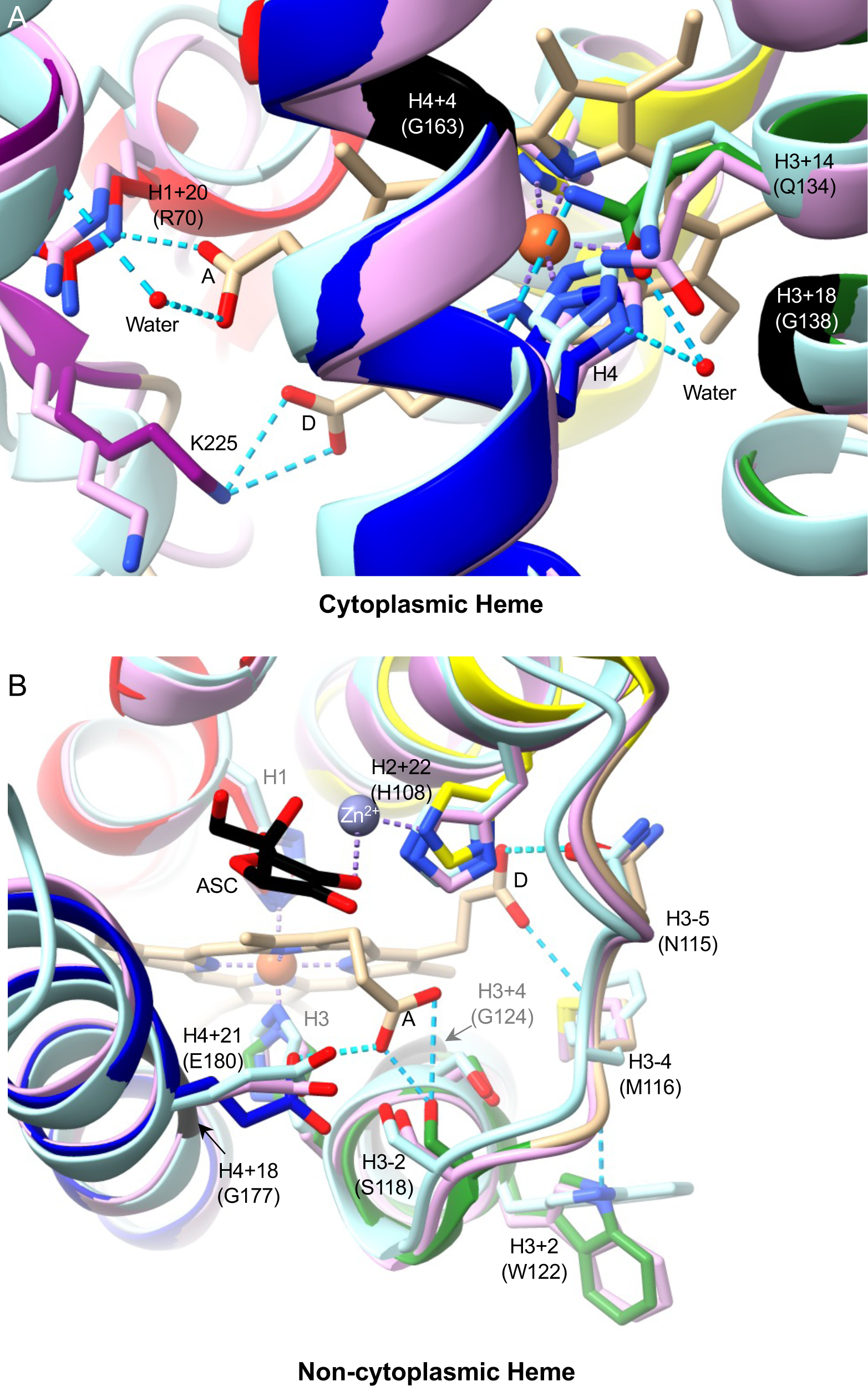
Heme binding by Dcytb and *D. melanogaster* CG1275 and Nemy. AlphaFold structures of *D. melanogaster* CG1275 and Nemy were overlayed with Dcytb (5ZLG) using Chimera Matchmaker. Dcytb loops are in tan and helices are shown in red (helix 2), yellow (helix 3), green (helix 4), dark blue (helix 5), and purple (helix 6). CG1275 is in pale pink, and Nemy is in pale blue. Ascorbate (ASC) in black stick, heme in tan stick (with iron shown as orange spheres), and zinc (Zn^2+^) ion coordination in purple are from the 5ZLG structure. Select side chains are shown in stick form and are labeled by position relative to conserved heme-coordinating histidines with Dcytb amino acid numbering in parentheses. Key glycines are shown in black ribbon. Stick side chains, ascorbate, and heme have oxygen in red and nitrogen in blue. Hydrogen bonds shown as cyan dashed lines, including to the A and D heme-propionates (labeled as in [18]). Water molecules shown as red spheres. A) Cytoplasmic heme region. B) Non-cytoplasmic heme region. Hydrogen bonds predicted by ChimeraX or observed in Dcytb structure (Asn115 to D heme-propionate; Glu180 to A heme-propionate, [18]).

In the Dcytb and AtCytb561-B structures, we observed interactions between H4 and the glutamine at H3+14 (Gln134). Gln134 forms a predicted hydrogen bond with the H4 backbone carbonyl and contributes to a network of hydrogen bonds linked by a water molecule to the H4 imidazole ring positioned on the opposite side of the heme iron coordination site (Fig 6A). An H3+14 glutamine residue is conserved in all CG1275, Nemy, and CG8399 sequences, suggesting a role in the positioning of H4 in these three protein groups. In Group 4 sequences, a serine (16 out of 25) or threonine (4 out of 25) is often present at H3+14 (S1 Fig) that could both potentially interact with H4, perhaps through hydrogen bonding mediated by water. Thus, within CG1275, Nemy, and CG8399 proteins, a glutamine at H3+14 may help position H4 to coordinate the heme, while a salt bridge may form between the cytoplasmic heme A propionate and the arginine at H1+20. In Group 4 proteins, polar residues at H1+20 and H3+14 could conceivably have similar heme positioning roles.

We found that two of the highly-conserved small residue positions in insect cytb561s are predicted to form cytoplasmic heme contacts in Dcytb: H3+18 (Gly138) and H4+4 (Gly163) (S3 Table). A glycine at H4+4 is conserved across all insect groups, in addition to more diverse cytb561s [3,19], while a glycine at H3+18 is conserved in all insect groups except CG8399, where an alanine substitution occurs (Fig 2). It is likely that small residues at H3+18 and H4+4 are important for proper cytoplasmic heme positioning within the protein’s central cavity in diverse cytb561s (Fig 6A).

#### Non-cytoplasmic heme

We performed a similar analysis of non-cytoplasmic heme positioning. In Dcytb and AtCytb561-B, two residues form hydrogen bonds with the non-cytoplasmic heme A-propionate group: a serine at H3-2 (Ser118) and a glutamate at H4+21 (Glu180) (Fig 6B)[13,18]. These residues are well conserved in CG1275 and Nemy sequences and form predicted contacts with the heme group in the *D. melanogaster* CG1275 and Nemy models (Fig 4; S3 Table). As mentioned above, it is notable that the most highly-conserved regions among CG1275 and Nemy proteins are the sequences surrounding H1 and H3, which coordinate the non-cytoplasmic heme group (Fig 2). While CG8399 and Group 4 proteins share less conservation in these regions, a serine or threonine at H3-2 is well conserved across Group 4 proteins, and CG8399 sequences most often contain an asparagine (Fig 2; S1 Fig); these polar residues could potentially hydrogen bond with the non-cytoplasmic heme A-propionate (H4+21 is slightly beyond the conserved core cytb561 domain and was therefore omitted from our analysis of CG8399 and Group 4 proteins).

In Dcytb and AtCytb561-B, an asparagine at H3-5 forms a hydrogen bond with the non-cytoplasmic heme D-propionate (Fig 6B) [13,18]. This position is highly conserved in CG1275 and Nemy proteins (Fig 4); two Nemy sequences have a histidine, which also could form a potential interaction with a heme propionate. All Group 4A and most Group 4B proteins also have a histidine at H3-5 (Fig 2; S1 Fig). In contrast to the other groups, there is no clear conservation at H3-5 in CG8399 proteins, which is part of the core domain non-cytoplasmic loop (Fig 2; S1 Fig).

Additional AtCytb561-B residues that form hydrogen bonds with the non-cytoplasmic heme D-propionate are the side chain of H2+22 (His106) and the amide nitrogen of H3-4 (Phe114) [13]. Interestingly, in Dcytb the histidine at H2+22 is conserved but was reported to coordinate with a non-cytoplasmic zinc ion (Fig 6B, also discussed in more detail below) [18]. All but two CG1275 and Nemy sequences have a histidine at H2+22, whereas this position is most commonly a lysine in Group 4 proteins or a hydrophobic residue in CG8399 proteins (Fig. 2; S1 Fig). Dcytb has a methionine (Met116) at H3-4 whose amide nitrogen is also positioned to hydrogen bond with the non-cytoplasmic heme D-propionate, similarly to Phe114 in AtCytb561-B (Fig 6B). In CG1275 and Nemy sequences, a methionine, phenylalanine, or leucine is found at H3-4 and a phenylalanine is also common in Group 4 proteins (18 out of 25) (Figs 2 and 4). In contrast, CG8399s have no notable conservation at this position (S1 Fig). The hydrophobic side chain at H3-4 in Dcytb (Met116) appears to anchor the non-cytoplasmic loop between helices 3 and 4 in the membrane, and its backbone carbonyl is predicted to form a hydrogen bond with the sidechain of the tryptophan (Trp122) at H3+2, providing added stability to the loop region between helices 3 and 4 (Fig 6B). A tryptophan at H3+2 is entirely conserved in CG1275 and Nemy proteins (Fig 4), suggesting a similar positioning of the non-cytoplasmic heme and loop between helices 3 and 4. The tryptophan at H3+2 is not conserved in CG8399 or in most Group 4 proteins (Fig 2).

Glycine residues at positions H3+4 (Gly124) and H4+18 (Gly177) are predicted to make non-cytoplasmic heme contacts in the Dcytb, CG1275, and Nemy structures (Figs 4 and 6B; S3 Table) These residues were previously identified as highly conserved across diverse cytb561s [3]. The H3+4 glycine is conserved in all CG1275, Nemy, and CG8399 sequences, and is highly conserved in Group 4 proteins (21 out of 25) (Fig 2). A glycine at H4+18 is conserved in all CG1275 and Nemy sequences and most Group 4 proteins (19 of 25) also have a glycine at H4+18, whereas CG8399s have an alanine substitution (Fig 2; S1 Fig). Therefore, it is likely that small residues at H3+4 and H4+18 (non-cytoplasmic side), as well as H4+4 and H3+18 (cytoplasmic side, described above) are important for positioning the bulky heme groups in the central cavity of the protein across a range of cytb561s.

#### Heme positioning summary

Overall, heme positioning appears to be similar in Dcytb, AtCytb561-B, CG1275, and Nemy proteins, whereas heme positioning in CG8399 and Group 4 proteins is less similar. Non-cytoplasmic heme positioning is most dissimilar in CG8399 proteins, and cytoplasmic heme positioning is most dissimilar in Group 4 proteins.

### Ascorbate binding

Dcytb and AtCytb561-B were found to bind to ascorbate (or the closely related monodehydroascorbate) in the cytoplasm through the interactions of basic residues with some of the ascorbate oxygens [13,18]. On the non-cytoplasmic side, phenylalanine was implicated in stacking with the ascorbate ring [13,18]. We located these Dcytb and AtCytb561-B residues in relation to the conserved heme-coordinating histidines and examined our insect sequences for conserved residues, especially basic and aromatic residues, at these locations. We also performed ChimeraX contact analysis with the Dcytb structure and overlayed CG1275 and Nemy structures to identify other amino acids proximate to ascorbate to identify similarities and differences in how ascorbate might bind.

#### Potential cytoplasmic ascorbate binding residues

The Dcytb and AtCytb561-B crystal structures clearly identified three basic residues that bind cytoplasmic ascorbate: H2-7 (Lys79), H2-3 (Lys83), and H4-7 (Arg152) [13,18]. Additionally, ChimeraX analysis predicted that Dcytb has ascorbate contacts at H3+22 (Phe142) and H4-3 (Met156) (Fig 7A; S3 Table). These five residues are also predicted to have cytoplasmic ascorbate contacts in *D. melanogaster* CG1275 (S3 Table), and most CG1275 proteins have these conserved residues except for some variation at H4-3 (Fig 4). In Nemy sequences, a lysine at H2-3, an arginine at H4-7, and a phenylalanine at H3+22 are entirely conserved, whereas either a valine or threonine is present at H4-3 (Fig 4); *D. melanogaster* Nemy is predicted to have ascorbate contacts at these four positions (S3 Table). More variation is seen at Nemy H2-7, which is most often an arginine (4 out of 8) and in one sequence is a lysine (Fig 4). However, H2-7 is not a basic residue in the *D. melanogaster*, *A. gambiae*, and *A. pisum* Nemy sequences (Fig 4). The *D. melanogaster* Nemy model suggests a shift in the placement of the lysine at H2-3 (Fig 7A), which may compensate for the lack of a basic residue at H2-7. Additionally, the Nemy model predicts that a serine (Ser205) at H4-6 hydrogen bonds with ascorbate, suggesting a unique functional role for this serine in *D. melanogaster* Nemy and in *A. gambiae* Nemy, which also has a serine at this position (Fig 7A).

**Fig 7.**
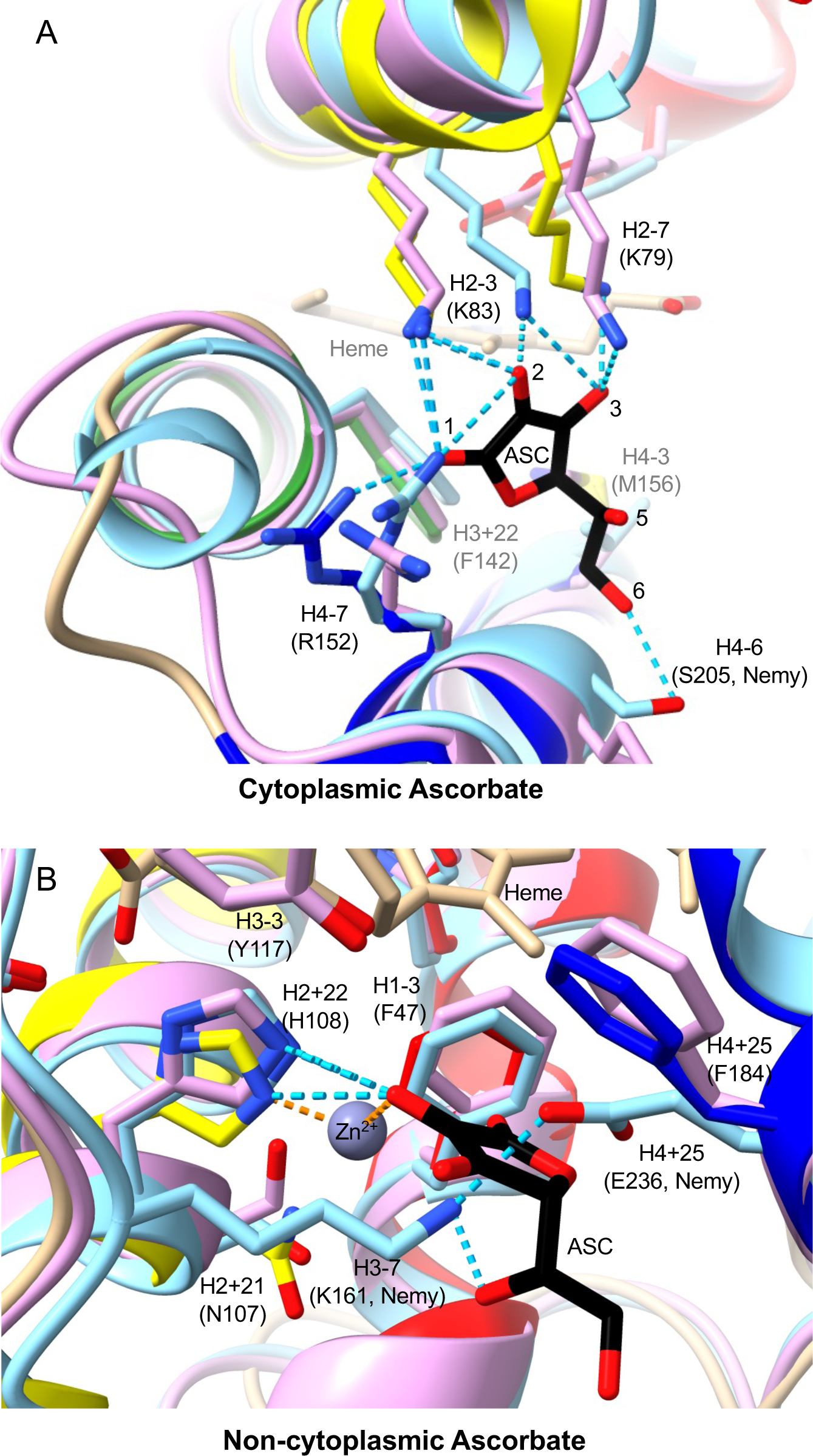
Ascorbate binding in Dcytb, CG1275, and Nemy. AlphaFold structures of *D. melanogaster* CG1275 and Nemy were overlayed with Dcytb (5ZLG) using Chimera Matchmaker. Dcytb loops are in tan and helices are shown in yellow (helix 3), green (helix 4), and dark blue (helix 5). CG1275 is in pale pink, and Nemy is in pale blue. Ascorbate (ASC) in black stick, heme in tan stick (with iron shown as orange spheres), and metal ion coordination in purple are from the 5ZLG structure. Select side chains are shown in stick form and are labeled by position relative to conserved heme-coordinating histidines with Dcytb amino acid numbering in parentheses (unless otherwise noted as Nemy numbering). Stick side chains, ascorbate, and heme have oxygen in red and nitrogen in blue. Hydrogen bonds shown as cyan dashed lines. A) Cytoplasmic ascorbate region. Ascorbate oxygens involved in bonding are labeled 1-3 and 6. B) Non-cytoplasmic ascorbate region with predicted hydrogen bonds to Dcytb, CG1275, and Nemy shown in cyan; an additional predicted hydrogen bond between Nemy Lys161 and Glu236 is also shown. Zinc coordination bonds are shown as orange dashed lines.

Of the five residues proximate to cytoplasmic ascorbate in Dcytb and AtCytb561-B (basic residues at H2-7, H2-3, and H4-7, aromatic at H3+22, bulky hydrophobic at H4-3), only an arginine at H4-7 (Arg509) is completely conserved across CG8399 sequences (amino acid numbering in this section corresponds with *D. melanogaster* CG8399) (Fig 2). Most CG8399 sequences also have a conserved lysine at H2-7, although *D. melanogaster* CG8399 has a threonine (Thr443) (S1 Fig). To visualize conservation on the cytoplasmic side of CG8399 proteins, we overlayed the CG8399 consensus sequence alignment onto the *D. melanogaster* CG8399 AlphaFold model (Fig 8). Of the remaining positions discussed in the interaction of cytoplasmic ascorbate with Dcytb, AtCytb561-B, CG1275, and Nemy proteins (H2-3, H3+22, and H4-3), CG8399 sequences have conserved residues with differing properties. At H2-3, CG8399s have a phenylalanine (Phe447) rather than the lysine found in Dcytb, AtCytb561-B, CG1275, and Nemy proteins, and a conserved tryptophan (Trp446) precedes H2-3 at H2-4 (Fig 8). Instead of a bulky hydrophobic residue at H3+22, a proline (Pro502) is present in CG8399s; that proline is preceded by a completely conserved CG8399 arginine (Arg501) at H3+21 (Fig 8). Instead of a bulky hydrophobic residue at H4-3, CG8399 proteins have an asparagine (Asn513). We noted additional highly-conserved positions in this region among CG8399 proteins, including a second proline at H3+24 (Pro504) and a basic residue at H1+23 (Lys432), which is either a lysine or arginine in all the CG8399 sequences. H1+23 is an arginine in most CG1275 and Nemy sequences although not in Dcytb (Trp73) or AtCytb561-B (Gln74) (Fig 4). While side-chain placement cannot be determined with high confidence, it is notable how highly conserved this region is across multi-domain CG8399 proteins, including conserved residues suitable for binding ascorbate: basic residues at H1+23, H3+21, and H4-7, and aromatics at H2-4 and H2-3 (Fig 8; S5 Fig).

**Fig 8.**
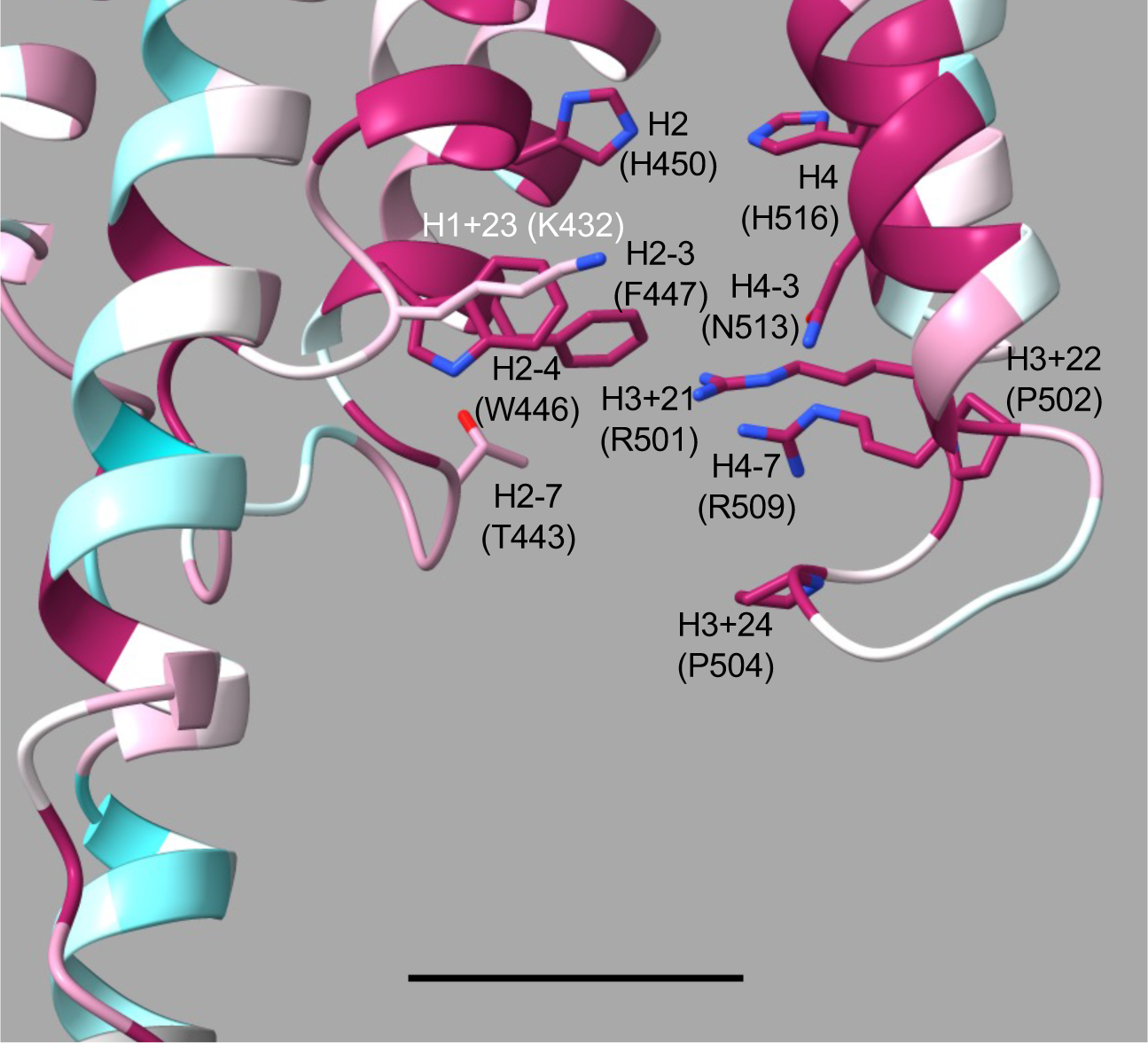
Cytoplasmic side of *D. melanogaster* CG8399 model. AlphaFold structure of *D. melanogaster* CG8399 overlayed with multi-domain CG8399 consensus sequence alignment. The model is gradient colored with high conservation (1.68 score) in magenta to low conservation (−1.89 score) in cyan. Select side chains are shown in stick form with oxygen in red and nitrogen in blue and are labeled by position relative to conserved heme-coordinating histidines with *D. melanogaster* CG8399 amino acid numbering in parentheses. Ascorbate binding on the cytoplasmic surface of CG8399 may be facilitated by a cluster of aromatic and basic residues: Lys432, Trp446, Phe447, Arg501, and Arg509. Also shown are conserved H2 (His450) and H4 (His516), which coordinate the cytoplasmic heme iron (not present in model). Two well-conserved prolines at positions H3+22 and H3+24 stabilize the loop between helices 3 and 4. Other cytoplasmic side positions with well conserved residues include H2-7, which typically has a lysine but is Thr443 in *D. melanogaster*, and H4-3 (Asp513). Scale bar depicts 10 Å.

Some Group 4 proteins also have a basic residue at two of the three positions that hydrogen bond with cytoplasmic ascorbate in Dcytb and AtCytb561-B: H2-7 (11 out of 25) and H4-7 (11 out of 25) (S1 Fig). While only six Group 4 proteins have a basic residue at H2-3, nearly all have basic residues at H2-4 (23 out of 25) and H4-3 (23 out of 25) (S1 Fig). This is notable since H4-3 is one of the two positions we identified as having a potential hydrophobic contact with cytoplasmic ascorbate in Dcytb, CG1275, and Nemy (Fig 7A; S3 Table). It is possible that the basic residues at these highly-conserved positions are involved in Group 4 cytoplasmic ascorbate binding. At the other position (H3+22) with predicted hydrophobic ascorbate contact, most Group 4 proteins (19 out of 25) also have a large, hydrophobic residue that could potentially interact with ascorbate or other possible cytoplasmic reducing agents, such as dihydrolipoic acid (S1 Fig)[53]. The adjacent position, H3+23, is also typically an aromatic residue (17 out of 25) or occasionally a lysine (3 out of 25) and may be involved in cytoplasmic ascorbate binding (S1 Fig).

#### Non-cytoplasmic interactions with ascorbate and zinc: implications for ferric reductase activity

Dcytb and AtCytb561-B also bind ascorbate at their non-cytoplasmic surfaces [13,18]. In Dcytb, this appears to be part of its ferric reduction mechanism [18]. To identify an iron-binding site in Dcytb, the protein was crystallized (PDB: 5ZLG) in the presence of redox inactive zinc, which presumably binds Dcytb in a manner similar to iron [18]. Details of Dcytb ascorbate-complexed metal binding are described below, and this information was used to predict non-cytoplasmic ascorbate and metal binding by insect cytb561 proteins.

In Dcytb the only residue found to directly interact with non-cytoplasmic ascorbate was Phe184 (H4+25) via a van der Waals interaction, although several residues were discussed as contributing to the environment of the ascorbate-binding pocket [18]; in AtCytb561-B the same H4+25 (Phe182) interaction was reported, and several residues were also suggested to form possible hydrogen bonds, including H2+22 (His106) and H3-3 (Tyr115) [13]. Our analysis revealed that three Dcytb residues have predicted contacts with non-cytoplasmic ascorbate: H1-3 (Phe47), H2+22 (His108), and H4+25 (Phe184) (Fig 7B; S3 Table). A phenylalanine at H1-3 is also predicted to make ascorbate contacts in *D. melanogaster* CG1275 and Nemy (S3 Table); a phenylalanine at this position is conserved across CG1275 and Nemy sequences, and is centrally located – near the zinc ion, the non-cytoplasmic ascorbate, and the non-cytoplasmic heme – in Dcytb, CG1275, and Nemy structures (Fig 7B). In Dcytb, Phe47 was the only residue that had an observed change in electron density upon zinc ion and ascorbate binding, indicating a change in position [18]. Most Group 4A proteins have a phenylalanine at H1-3; however, there is less conservation at this position in CG8399 and Group 4B proteins (S1 Fig).

As mentioned above, the phenylalanine (Phe184) at H4+25 was determined to form a van der Waals interaction with the non-cytoplasmic ascorbate in Dcytb and AtCytb561-B [13,18]. An aromatic amino acid, most often phenylalanine, is typical at H4+25 across CG1275 and Nemy sequences with a small number of exceptions; the most dramatically different is a glutamate (Glu236) in *D. melanogaster* Nemy (Fig 4). In the absence of ascorbate, our analysis revealed that Glu236 and a lysine at H3-7 (Lys161) are predicted to form a hydrogen bond in the *D. melanogaster* Nemy model that would run straight through the region occupied by the non-cytoplasmic ascorbate ring (Fig 7B). The *D. melanogaster* Nemy model also has extensive contacts predicted between the non-cytoplasmic ascorbate and Lys161 (23, including one hydrogen bond) and Glu236 (29) (S3 Table). These results suggest that *D. melanogaster* Nemy could bind non-cytoplasmic ascorbate though a combination of compensatory substitutions in lieu of an aromatic residue at H4+25. H4+25 is slightly outside the conserved core domain and therefore was not included in our analyses of CG8399 and Group 4 sequences.

In Dcytb, the histidine (His108) at H2+22 was determined to be part of an ascorbate-metal-binding complex; His108 was found to directly coordinate with a redox inactive zinc ion, which directly interacted with the non-cytoplasmic ascorbate [18]. The histidine at H2+22 is conserved across CG1275 and Nemy sequences and is predicted to be similarly positioned in *D. melanogaster* CG1275 and Nemy models (Fig 4; Fig 7B). In contrast, a histidine at H2+22 is not conserved in other insect cytb561s (Fig 2). The residue at H2+22 is nonpolar in CG8399 sequences, whereas many Group 4 proteins have a lysine at this position, which could contribute to ascorbate binding but would not be expected to coordinate a metal cation (Fig 2; S1 Fig).

His108 was also found to be critical for Dcytb ferric reductase activity, as were residues at H2+21 (Asn107), H3-3 (Tyr117), and H4+25 (Phe184, discussed above) [18]. Like Dcytb, CG1275 and Nemy proteins have a tyrosine at H3-3 in addition to the histidine at H2+22 and phenylalanine at H4+25, and they tend to have a conservatively-substituted serine at the H2+21 position (Fig 2). These conserved residues support the hypothesis that CG1275 and Nemy proteins have ferric reductase activity. In contrast, conservation of these residues in the other insect cytb561 proteins is limited to a glutamate that is entirely conserved at H2+21 in multi-domain CG8399 proteins (discussed further below) (Fig 2).

Some Group 4A proteins may have a different type of non-cytoplasmic ascorbate-binding site. A lysine-containing “KXXXXKXH” motif was observed in the non-cytoplasmic loop of the core domain in six of the seven Group 4A proteins (excluding one of the two *C. felis* sequences) (Fig 2; S1 Fig). Group 4A includes the proteins identified as more like human TScytb than any *D. melanogaster* cytb561 (Table 1; S2 Table). We aligned these six Group 4A proteins with human TScytb (which can reduce ferric ions in nanodiscs [14,54]) and TScytb-like proteins of diverse animal species and found that all sequences have this “KXXXXKXH” motif (Fig 9A), which contributes to the high degree of conservation across TScytb proteins on the non-cytoplasmic side (Fig 9B). The location of the “KXXXXKXH” motif in the non-cytoplasmic loop between helices 3 and 4 (where substrate binding occurs in Dyctb), as well as its strict conservation in TScytb-like proteins of diverse species, suggests an important functional role, possibly related to ascorbate binding (Fig 9C). This motif was not identified in other human or insect cytb561s. A completely conserved tryptophan at H4+23 could also potentially contribute to a non-cytoplasmic binding site, and the histidine at H3-5 (within the “KXXXXKXH” motif) could potentially coordinate a ferric ion complexed with ascorbate, similar to the substrate binding site of Dytcb, providing a site for ferric reduction in TScytb-like proteins (Fig 9C).

**Fig 9.**
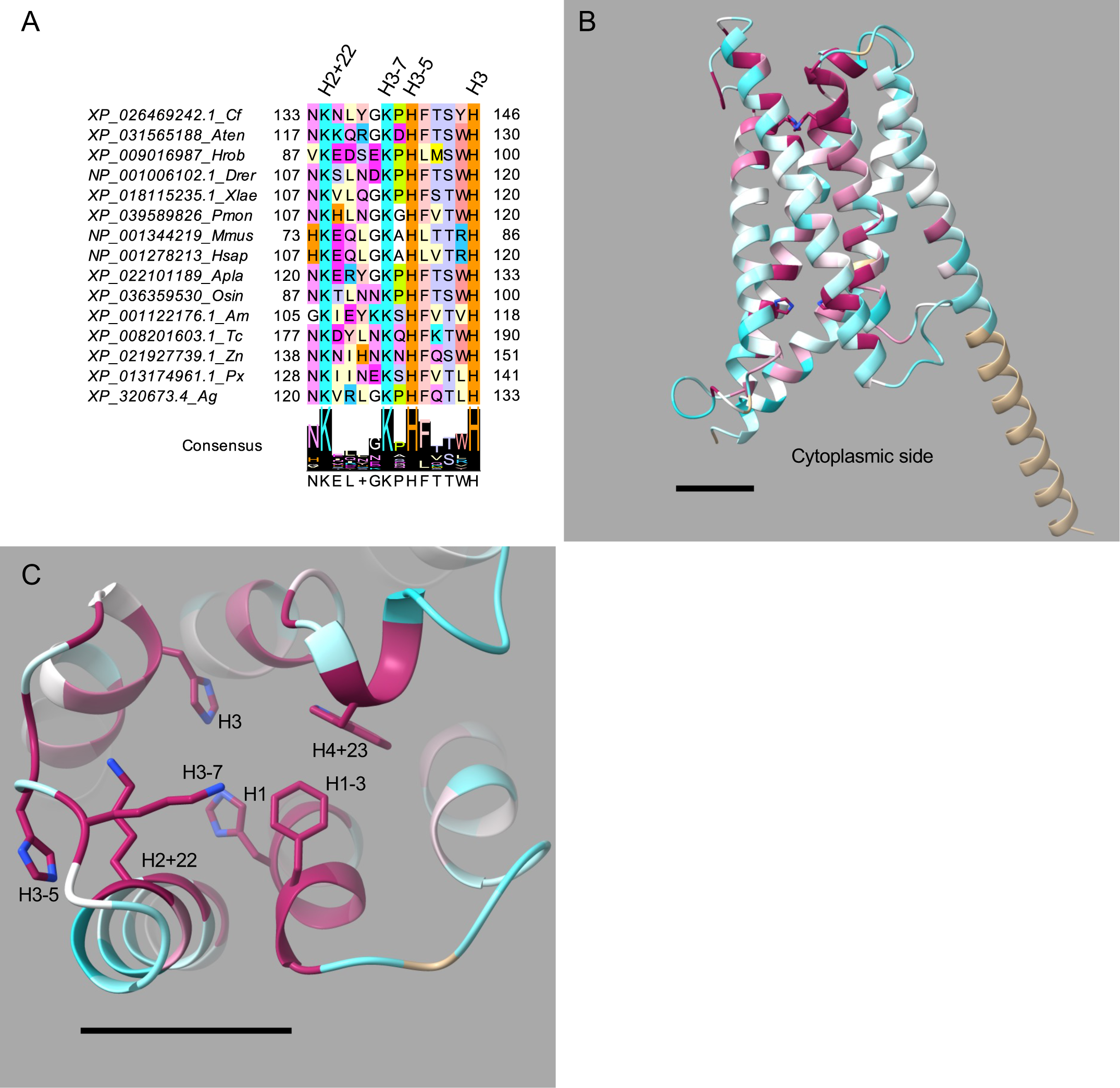
A conserved “KXXXXKXH” motif in Group 4A and related proteins. A) Truncated alignment of animal TScytb and Group 4A sequences shows a conserved “KXXXXKXH” motif upstream of the conserved non-cytoplasmic heme-coordinating H3. Coloring by amino acid residue as in Fig 4. Abbreviations: Cf, *C. felis*; Aten, *A. tenebrosa*; Hrob, *H. robusta*; Drer, *D. rerio*; Xlae, *X. laevis*; Pmon, *P. montanus*; Mmus, *M. musculus*; Hsap, *H. sapiens*; Apla, *A. planci*; Osin, *O. sinensis*; Am, *A. mellifera*; Tc, *T. castaneum*; Zn, *Z. nevadensis*; Px, *P. xuthus*; Ag, *A. gambiae*. B) Full alignment of animal Tscytb and Group 4A sequences overlayed on AlphaFold model of *A. gambiae* XP_3200673.4 (AF-Q7PZD0-F1-model_v4). H1-H4 shown in stick form with nitrogen in blue. C) Non-cytoplasmic side of the protein contains completely conserved lysines (H2+22, H3-7), a histidine (H3-5), a tryptophan (H4+23), and a phenylalanine (H1-3), in addition to H1 and H3; these residues are shown in stick with nitrogen atoms in blue. B-C) Amino acid conservation is shown by a gradient of color as in Fig 3. Scale bars show 10 Å.

### Other amino acids of interest

Aside from H1-H4, the most highly-conserved residues in the core domain of insect cytb561s are small residues and hydrophobic residues (S1 Fig). As discussed above, four well-conserved, small residues (H3+4, H3+18, H4+4, and H4+18) appear to be important for heme placement. The remaining well-conserved residues may play a role in helical packing [55]. The small amino acids at H1+7 and H2+14 are positioned where TM2 and TM3 face each other, while the small residue at H1+13 is on the side of TM2 that faces TM5.

Of the investigated insect cytb561s, Nemy proteins possess unique cysteine-rich sequences in their cytoplasmic loops, with double cysteines common in both cytoplasmic loops of the core domain (S1 Fig). This observation is notable, as cysteines are among the least common amino acids in proteins, yet they are often at positions of functional importance [56]. Cytoplasmic cysteine residues can participate in redox-active disulfide formation and metal ligation, and they can also serve as sites of post-translational modifications [56]. While the pH and reducing environment of the cytoplasm is not conducive to disulfide bond formation, solvent exposed cysteines can be regulated by redox state [57]. The cytoplasm-exposed Nemy cysteines could form redox active disulfides, potentially functioning as an on/off switch for electron transfer activity. Another possibility involves cysteine S-palmitoylation, given that *D. melanogaster* Nemy was identified as a palmitoylated protein [58]. Cysteine S-palmitoylation is a reversible post-translational modification used to regulate protein function [59]. Given its dynamic nature, cysteine S-palmitoylation can direct varied cellular processes, including subcellular trafficking, protein localization, and enzyme activity [59]. We did not detect multiple cysteine-rich cytoplasmic loop sequences in the cytb561 domain of other insect cytb561s, nor in human cytb561 proteins, suggesting a function unique to Nemy proteins. Future research could indicate whether these cysteines are palmitoylated and whether they influence Nemy localization, cytoplasmic substrate binding, or protein function.

### Analysis of the CG8399 DOMON domain

#### Lack of conservation in the extracellular sections of the CG8399 cytb561 domain suggests DOMON domain involvement in ferric reductase activity

As discussed above, many of the Dyctb non-cytoplasmic surface residues that interact with ascorbate or function in ferric reduction are missing in CG8399 proteins (Fig 2) [13,18]. Nevertheless, *D. melanogaster* CG8399 and its ortholog in mice, SDR2, possess in vitro ferric reductase activity [6,31]. These observations indicate that ferric reductase activity in CG8399 and SDR2 occurs through a mechanism different from that of Dcytb and suggests the possibility that iron-binding in CG8399 and SDR2 occurs in the extracellular DOMON domain. It has been previously proposed that CYBDOMs may act as a “redox bridge” between cytosolic ascorbate and the extracellular environment [1]. Using the redox bridge concept as a model, we suggest that the CG8399 cytb561 domain binds the reducing agent (ascorbate) on the cytoplasmic surface, then transfers the electron to a DOMON domain heme, which then transfers the electron to an electron-accepting substrate (possibly a metal ion).

#### CG8399 DOMONs may bind a b-type heme

To further investigate the DOMON domain in CG8399 proteins, we used the HHpred PDB server to identify proteins with known structure that have the highest probability percentage of homology to CG8399 DOMON sequences. We found that all insect CG8399 DOMONs have > 99% probability of homology to the DOMON domain of human DBH and the cytochrome domain of *P. chrysosporium* CDH. (Note that despite the use of “cytochrome” in the domain name of *P. chrysosporium* CDH, this domain is not homologous with the cytochrome b561 domain of CG8399 proteins; rather, the *P. chrysosporium* CDH cytochrome domain is homologous with the DOMON domain in CG8399 proteins.) Intriguingly, DBH and CDH have only 10% sequence identity and bind different ligands, yet their structural alignment suggests that both domains have very similar folds [60].

The crystal structure of the CDH cytochrome domain (DOMON homolog) revealed a b-type heme group coordinated by an unusual methionine and histidine ligation of the heme iron [61]. HHpred alignments of CDH with each individual insect CG8399 DOMON showed conservation of these methionine and histidine residues in all sequences (Met286 and His379 in *D. melanogaster* CG8399 [S2 Fig]). Notably, the methionine and histidine are not conserved in the DOMONs of proteins that bind sugar rather than heme, nor in DBH, which binds neither sugar nor heme [20,60]. We then generated an alignment of CG8399 DOMON domains with the DOMON domains of diverse animal CYBDOM proteins, including insect CG8399s, and found universal conservation of the methionine and histidine across all species investigated, including both DOMON domains in the *A. pisum* CG8399 (S3 Fig). These results agree with previous observations that CYBDOM proteins likely have a b-type heme group in their DOMON domain [1]. Given the DOMON domain’s suspected role in interdomain electron transfer and the likelihood that CG8399s and their CYBDOM orthologs coordinate a heme group within their DOMON domains, it seems likely that their putative ferric reductase activity occurs outside of the cytb561 domain.

#### Spatial relationship between the DOMON domain and cytb561 domain in CG8399

To study the possibility of electron transfer to the DOMON domain, we analyzed the multi-domain CG8399 consensus sequence overlay of the AlphaFold model for D. *melanogaster* CG8399, which includes the extracellular reeler and DOMON domains in addition to the transmembrane cytb561 domain (Fig 10). In the following discussion, residue numbers relate to the *D. melanogaster* CG8399 protein sequence. The model predicts a DOMON domain largely composed of two five-strand beta sheets, similar to the DOMON domain in DBH [60]. The predicted aligned error plots for the DOMON and cytb561 domains were relatively low, suggesting their models could be informative for positional insights [47,48]. (In contrast, the predicted position of the reeler domain was reported with high error.) Notably, the *D. melanogaster* DOMON domain is oriented adjacent to the cytb561 domain, with predicted hydrogen bonds between the side chains of Lys285 and Arg383 in the DOMON domain with the backbone carbonyl oxygens of Leu472 and Pro538 in the cytb561 domain, respectively (Fig 10B). Salt bridges are predicted to form in the non-cytoplasmic loops of the cytb561 domain between Arg403 and Glu541, as well as Glu471 and Lys539 (Fig 10B).

**Fig 10.**
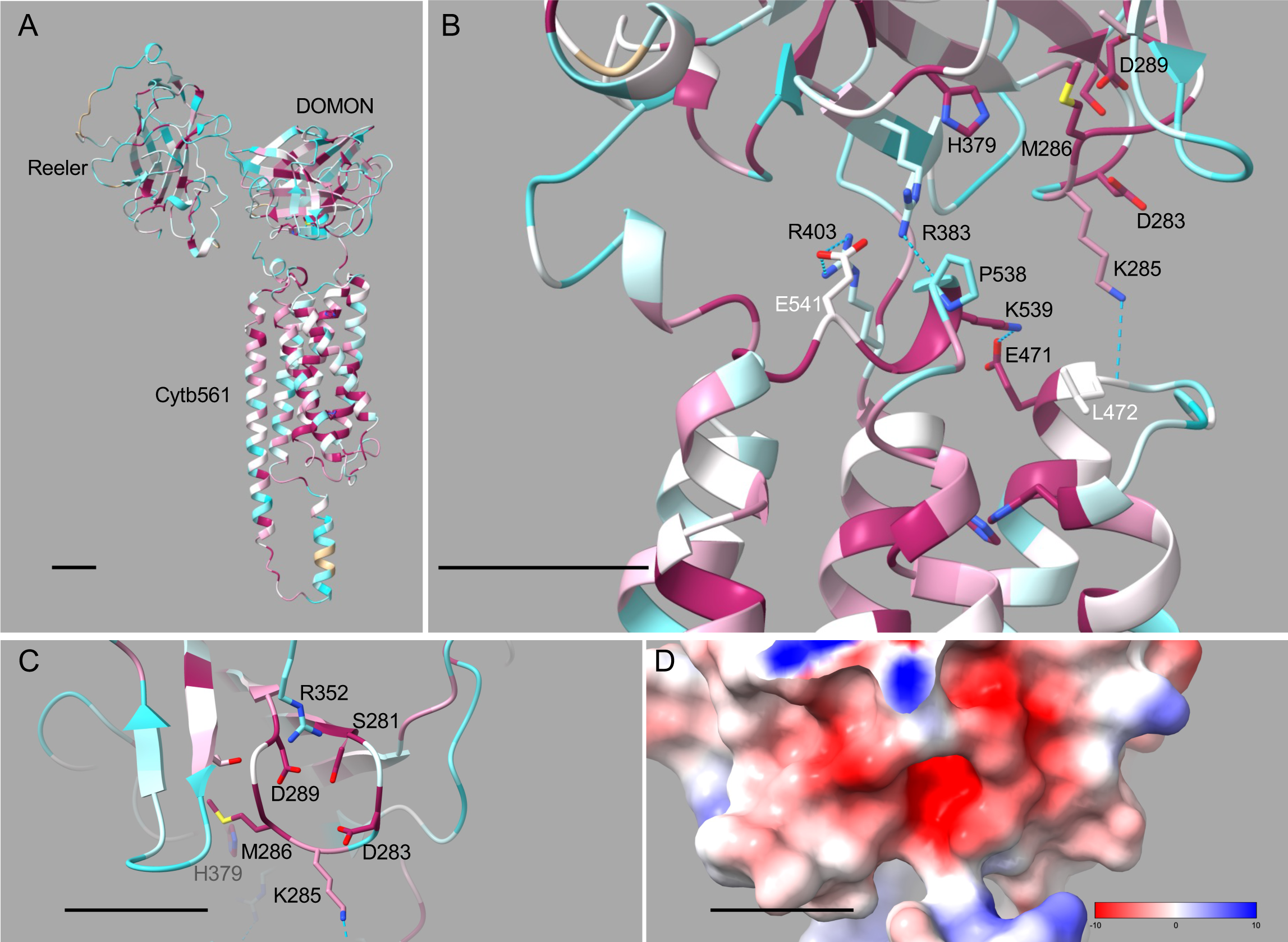
Model of *D. melanogaster* CG8399 demonstrating interactions between domains, and a putative iron binding site. A-C) Models were colored by ChimeraX based on amino acid conservation in the nine multidomain CG8399 sequences. Color gradient as in Fig 3. Residues in stick form show oxygen in red, nitrogen in blue and sulfur in yellow. A-D) Scale bars depict 10 Å. A) CG8399 has three distinct domains: reeler (top left), DOMON (middle right), and cytb561 (bottom). Residues expected to be heme-coordinating are shown in stick (some not visible). B) Selected sidechains at the interface of the DOMON (top) and cytb561 (bottom) domains. Predicted heme-coordinating residues (Met286 and His379) in the DOMON domain are positioned at the bottom of the domain where it is predicted to interact with the cytb561 domain. Interdomain hydrogen bonds were predicted between the sidechain of Lys285 (DOMON) and the backbone of Leu472 (cytb561), as well as the sidechain of Arg383 (DOMON) and the backbone of Pro538 (cytb561). Potential salt bridge interactions may form in the cytb561 domain loop regions between Glu471 and Lys539, as well as Arg403 and Glu541. C) A putative metal binding site in a highly conserved loop with two aspartates and a serine residue. D) Space filling model of the same part of the structure depicted in C, with Coulombic coloring calculated by ChimeraX, red for negative charge, white for neutral, and dark blue for positive charge.

We then located the predicted positions of the putative heme-coordinating residues (Met286 and His379) in the DOMON domain, with particular interest in their relative positions to the non-cytoplasmic heme-coordinating histidines (His409 and His480) in the cytb561 domain. The location of the putative DOMON heme-binding pocket is predicted to be directly adjacent to the non-cytoplasmic heme-binding region of the cytb561 domain (Fig 10B) [47,48]. These predictions further support the hypothesis that the CG8399 cytb561 domain functions in interdomain electron transport rather than being the site for oxidoreductase activity.

#### The CG8399 DOMON domain may contain a metal-binding pocket

If CG8399 ferric reductase activity occurs within the DOMON domain, we would expect to locate an iron-binding pocket within this domain. Our DOMON alignment revealed multiple well-conserved aspartates, including strict conservation of Asp283 and Asp289 (S3 Fig). Asp283 and Asp289 are in a highly-conserved region (sequence “SXDXXMGXD”) that includes the putative heme-coordinating methionine (Met286) (S3 Fig). This region also contains additional aspartates, especially in insect CG8399s, of which two are particularly well conserved (S3 Fig). We located this region in the *D. melanogaster* CG8399 model and found that a negatively-charged pocket is predicted to be directly behind the putative heme-binding pocket, similar to the position of the putative metal ion-binding pocket in DBH relative to its ligand-binding pocket (Figs 10C and D) [60]. These results are consistent with the DOMON domain being the site of ferric reductase activity, wherein strictly-conserved Ser281, Asp283, and Asp289 could form an iron-binding pocket in insect CG8399s and CYBDOMs from other species (S3 Fig). Metal-binding sites formed by aspartate residues have been suggested for other proteins. For example, the DBH DOMON domain has a putative metal ion-binding site composed of aspartate sidechains and the backbone carbonyl oxygens of two other amino acids [60], and three aspartate residues form a binding site for a variety of cations, including Fe^2+^ and Fe^3+^, in LarE [62]. This model also provides a possible explanation for the ferric reductase activity of *D. melanogaster* CG8399 and mouse SDR2, despite their lack of conserved ascorbate- and metal-binding residues on the non-cytoplasmic surface of the cytb561 domain.

### Three insect cytb561s with distinctive features outside of the core domain

Of the 54 identified insect cytb561s, we noted that three sequences possess unique features within the cytb561 domain but outside the domain core (TM2-TM5 in Nemy proteins, and TM1-TM4 in CG8399s). To determine whether other insect cytb561 proteins have any of these unique features, we performed additional BLAST searches against the entire class Insecta.

Compared to other Nemys, the *A. pisum* sequence has approximately 150 additional N-terminal residues preceding helix 1 and approximately 50 additional residues in the non-cytoplasmic loop connecting helix 1 and helix 2, which immediately precedes the core cytb561 domain (S4 Fig). We found proteins with similar extended regions in eight additional species of the Aphididae family, but not in other insects, suggesting that these regions may serve a unique function within aphids (S4 Fig; S5 Table). The extended N-terminal region preceding helix 1 is acidic and includes relatively high percentages of aspartate, glycine, serine, and threonine (S4 Fig).

Of the investigated CG8399s, only the *A. pisum* sequence possesses two DOMON domains. We identified 888 CYBDOM hits in the class Insecta, of which only 19 were found to have two DOMON domains: nine from other species of aphids, eight from other insects from the order Hemiptera, and two from insects in the order Coleoptera (S6 Table). It appears that multiple DOMONs are uncommon in insect cytb561s; however, they have a relatively high occurrence in Hemipterans. Both *A. pisum* DOMON domains contain the conserved methionine and histidine that are assumed to coordinate a heme group (S3 Fig); however, it is unknown how these two DOMON domains are spatially arranged.

While all nine investigated insect species possess a multidomain CG8399, *A. gambiae* (mosquito) is the only species with a second, single domain CG8399-like protein (XP_314065.2); this sequence lacks both reeler and DOMON domains. Further investigation revealed similar single-domain (cytb561 only) CG8399-like proteins in ten additional mosquito species, as well as two species of fungus gnats, which are also in the order Diptera (S7 Table). The single-domain CG8399-like protein is more similar to the cytb561 domain in multidomain CG8399 proteins than to the cytb561 domain in other groups of cytb561s (Table 1; S2 Table), including its core domain spanning TM1-TM4, rather than TM2-TM5.

### Cyb561 subcellular localization

Subcellular localization is important for protein function in an organism. We submitted each of the 54 full-length insect cytb561s to the SignalP-5.0 webserver for detection of predicted signal peptides, and to the DEEPLOC-1.0 webserver for prediction of subcellular protein localization [40]. Only the multidomain CG8399 proteins had a predicted signal peptide, and they are the only sequences with predicted plasma membrane localization (S8 Table). This result is consistent with the presence of *D. melanogaster* CG8399 in a plasma membrane fraction of fly head proteins [63]. The *A. gambiae* single-domain CG8399-like protein is predicted to be in the lysosome, as are all the CG1275 and Nemy proteins (S8 Table). The subcellular localization predictions for Group 4 proteins were not particularly robust, and the predicted location varied depending on the specific Group 4 protein (S8 Table).

An alignment of the Dcytb, Nemy, and CG1275 carboyxl-termini revealed that only the CG1275 proteins have a “EDXXLL” motif near TM6 that is consistent with a dileucine-based late-endosomal/lysosomal targeting signal “(D/E)XXXL(L/I)” (Fig 11) [42]. *A. mellifera* and *T. castaneum* CG1275 proteins lack one conserved leucine, and one of the four *A. pisum* sequences (XP_003246890.1) lacks both leucines, yet these three sequences are still predicted to localize to the lysosome (Fig 11; S8 Table).

**Fig 11.**
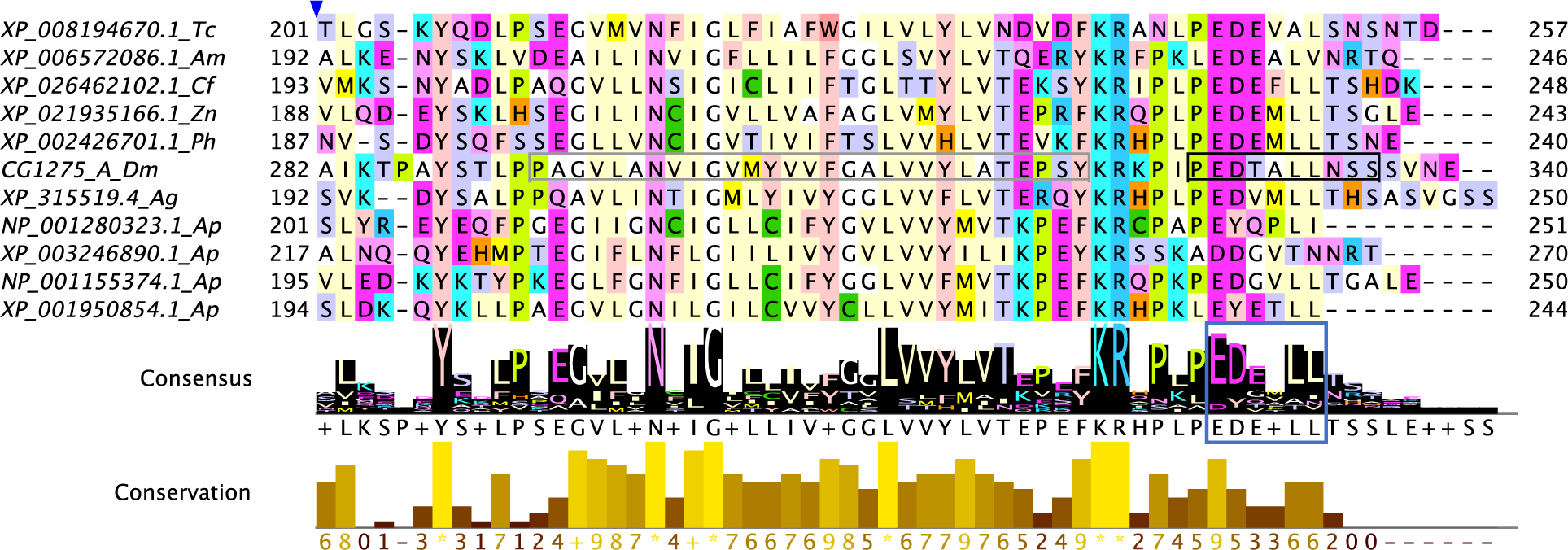
Lysosomal targeting signal in CG1275. An alignment of the C-terminal portion of CG1275 proteins. Helix predictions from the AlphaFold model of *D. melanogaster* CG1275 shown in gray (helix 6) and an additional helical region is in black. The EDXXLL motif is boxed in blue in the consensus logo.

### Summary of cytochrome b561s as potential ferric reductases in insects

A major goal of this study was to predict which of the insect cytb561 orthologous groups may function as ferric reductases and, thus, have a possible role in iron uptake by insect cells. Our analyses suggest that none of the four groups of insect cytb561 proteins can be excluded as ferric reductase candidates.

Based on sequence similarity, predicted structural similarity, and conservation of key amino acid residues, we conclude that CG1275 and Nemy proteins are strikingly similar to the well-established ferric reductase Dcytb, as well as the closely-related human proteins Lcytb and CGcytb. These similarities extend to the conservation of a metal-binding histidine and other amino acid residues that are required for ferric reductase activity by Dcytb. To date, no functions of CG1275 proteins have been discovered; however, if CG1275 proteins have ferric reductase activity, the constitutive, nearly ubiquitous expression of *D. melanogaster* CG1275 [64,65] would be consistent with a role in iron uptake by many cell types. We found that CG1275 proteins have a C-terminal dileucine motif that directs proteins to a late-endosomal/lysosomal location, suggesting that CG1275 proteins could function similarly to the non-homologous STEAP3 protein, which reduces Fe^3+^ to Fe^2+^ prior to uptake across endosomal and lysosomal membranes [27,28,66]. Among mammalian cytb561s, a C-terminal dileucine motif is present only in Lcytb proteins [7]. Interestingly, Lcytb appears to function as a lysosomal ferric reductase in Burkitt lymphoma cells, filling the role that STEAP3 plays in some other cell types [67]. Although Nemy proteins have many similarities with Dcytb and Lyctb, their one known function is unrelated to iron homeostasis [4,30]; however, Nemy proteins could be multifunctional enzymes that also play a role in iron uptake. *D. melanogaster* CG8399 and the mammalian ortholog SDR2 have in vitro ferric reductase activity [6,15,31], but the in vivo functions of CG8399 and SDR2 proteins are not known. Based on a previous biochemical study [63] and our subcellular localization predictions, CG8399 proteins localize to the plasma membrane, with their reeler and DOMON domains in the extracellular compartment. Their presence in the plasma membrane may enable reduction of extracellular ferric ions prior to iron uptake. Our analyses of the predicted structures of the CG8399 DOMON and cytb561 domains suggest that the two domains may function together in ferric reduction, with the interdomain transfer of an electron from cytoplasmic ascorbate to a heme in the extracellular DOMON domain and then to a ferric ion that is perhaps coordinated nearby by multiple aspartate residues. Because *D. melanogaster* CG8399 expression is limited to a subset of tissues [64,65], its hypothetical role in iron uptake would be limited to specific cell types.

We have the least amount of evidence to support the hypothesis that the diverse Group 4 proteins function as ferric reductases, and a lack of one-to-one orthologs indicates that these proteins may have non-conserved physiological functions. The expression profiles of all but one of the *D. melanogaster* Group 4 genes is moderately to highly restricted [64,65], excluding a role in iron uptake by most cell types. The Group 4 proteins have low similarity to the well-studied mammalian cytb561s, lacking even conserved residues that bind ascorbate in the cytoplasm. However, multiple basic residues and a large hydrophobic residue at the cytoplasmic surface of most Group 4 proteins may enable the Group 4 proteins to use ascorbate as an electron donor. On the non-cytoplasmic side, a subset of Group 4 proteins contain a “KXXXXKXH” motif that is also present in TScytb, a protein with an unknown biochemical function in vivo, but ferric reductase activity in vitro [14]. We propose that the “KXXXXKXH” motif may bind ascorbate and a ferric ion in a hypothetical ferric reduction site.

## Acknowledgements

We wish to acknowledge the undergraduate research students who participated in aspects of this project: Clare Burnett, Chrissy Chan, Caleb Garris, Keyata Lewis, Gregory Kane, Karen Nunez Sifuentes, Connor Sullivan, and Nataliya Yusopov. Thanks also to the MSU Denver CHE 4320 students from Spring 2018, Fall 2018, Spring 2019, Fall 2019 and Spring 2020 who participated in iterations of the Iron Uptake in Insects CURE. We thank Dr. Andrew McMillan for thoughtful comments on the manuscript. Molecular graphics and analyses were performed with UCSF ChimeraX, developed by the Resource for Biocomputing, Visualization, and Informatics at the University of California, San Francisco. FlyBase was used for gene identification and as a source of expression data.

