## Supplemental Figures 1-4 for "Comparison of insect and human cytochrome b561 proteins: Insights into candidate ferric reductases in insects"

**S1 Fig. Alignment of the homologous cytb561 core domain in all analyzed insect sequences.**

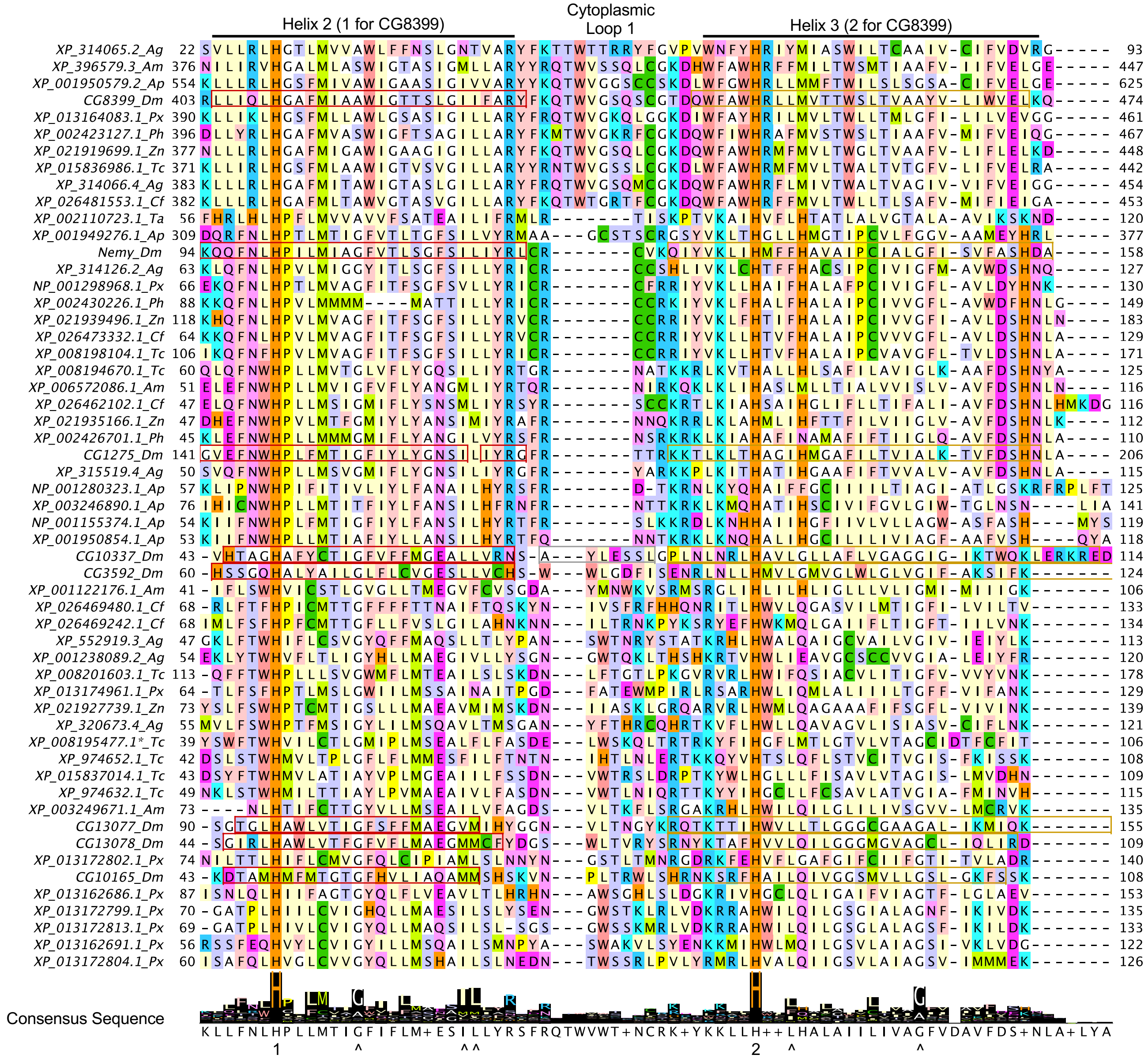

### S1 Fig. continued

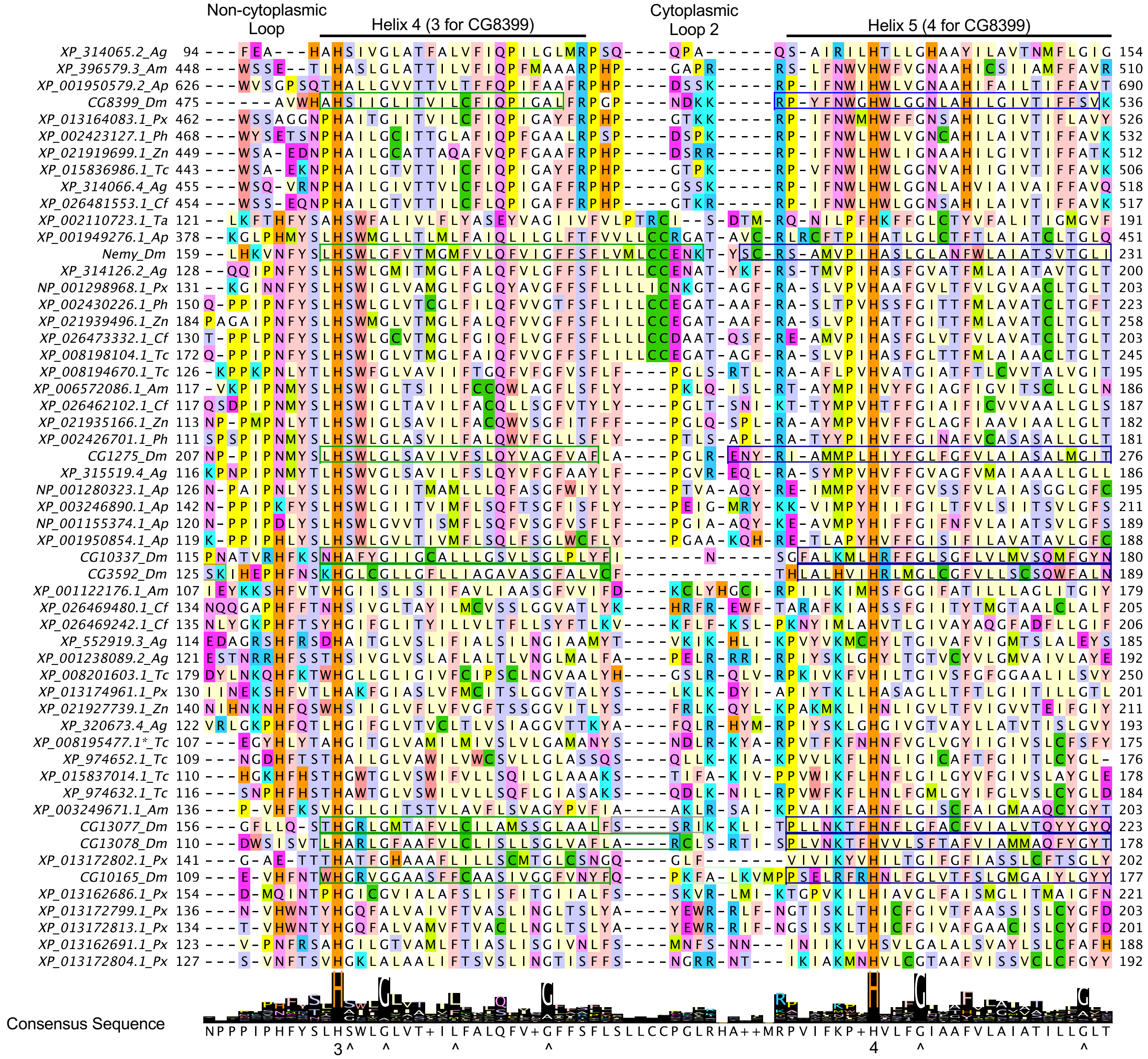

**S2 Fig. HHpred alignment between *D. melanogaster* CG8399 DOMON domain and *P. chrysosporium* cellobiose dehydrogenase (CDH) cytochrome domain (PDB 1D7B).**

```
                                                                *
1D7B_CDB_Pchr   24 DVTYGFVFPPLATSGAQSTEFIGEVVAPI----ASKW-IGIALG--GAMNNDLLLVAWA  75
CG8399_Dm      249 TSITVVTVRGD--------VFEFEIQSGKG--TNAA-YVAVGLSDDAKMGDDLTTECVP 296

1D7B_CDB_Pchr   76 NGN----QIVSSTRWAT-GY--V-QPTAYTGTATLTTLPETTINSTHWKWVFRCQGCTE 126
CG8399_Dm      297 ENG----RVNLYSSLTSASP-YSAVRSNVNQ-NSARLL-DASIVDGVIYCRVQRDAVTN 348

                                                         *
1D7B_CDB_Pchr  127 WNNGGGIDVTSQGVLAWAFSNVAVDDPSDPQSTFSEHTD---FGFFGIDYSTAHSAN 180
CG8399_Dm      349 VQGRTFDLRNGKYHLLVASGSSLKE------NSVGYHDIGRLPSAQPINLAEVQDLS 399
```

**S3 Fig. Alignment of DOMON domains from insect CG8399 proteins and CYBDOMs from other diverse species.**

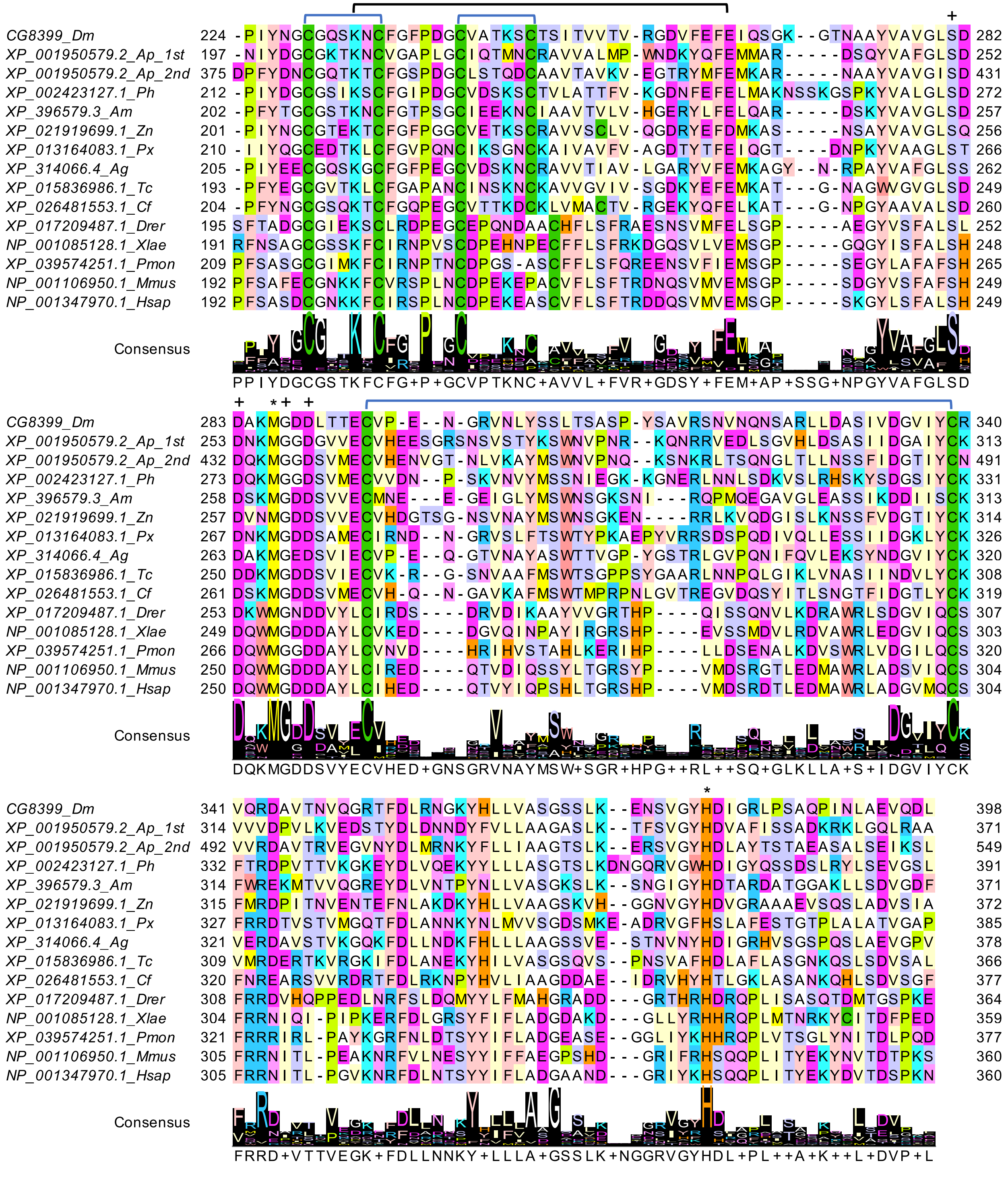

### S4 Fig. Alignment showing extended regions in the Aphid Nemy sequences

```
Nemy_A_Dm              ------------------------------------------------------------
VVC36571.1_Cc          1   ----MDSQQNTFLPPPIDGGEASAPLAQPDDDFVTANSAAGTPNTPADDAGTGSTVSPVS  56
KAE9533855.1_Agly      1   MDNQNNS-TTTFLPPPIDG-EASAPLAQPDDDFVVPLSAAGTPNTPADDGGTGSTVSPVT  58
XP_027844671.1_Ago     1   MDNQNNSTTSTFLPPPIDG-EASAPLAQPDDDFVVPLSAAGTPNTPADDGGTGSTVSPVT  59
KAF0754846.1_Ac        1   MDNQNNSNTTTFLPPPIDG-EASAPLAQPDDDFVVPLSAAGTPNTPADDGGTGSTVSPVT  59
XP_026816157.1_Rm      1   MDNQNNSTTTTFLPPPIDG-EASAPLAQPDDDFVVPLSAAGTPNTPADDGGTGSTVSPVT  59
XP_025196103.1_Ms      1   MDNQNNSTTSTFLPPPIDG-EASAPLAQPDDDFVVPLSAAGTPNTPADDGGTGSTVSPVT  59
XP_001949276.1_Ap      1   MDNQNNSTTTTFLPPPIDG-EASAPLAQPDDDFVVPLSVAGTPNTPADDGGTGSTVSPVS  59
XP_022171233.1_Mp      1   MDNQNNSNTTTFLPPPIDG-EASAPLAQPDDDFVVPLSAAGTPNTPADDGGTGSTVSPVS  59
XP_015366894.1_Dn      1   MDNQNNSTTTTFLPPPIDG-EASAPLAQPDDDFVVPLSAAGTPNTPADDGGTGSTVSPVS  59

Nemy_A_Dm              ------------------------------------------------------------
VVC36571.1_Cc          57  PVHLNRQKINAMMIDDDQVE-ADNDNGATDDSAGLLPPPDETDTDDKSTKDTSTTVTKND 115
KAE9533855.1_Agly      59  PVHLNRQPHAL-MMDDDQAANAATDSGATDDNSVLLPPPEETDSEEKPNKDGGDDGDNGG 117
XP_027844671.1_Ago     60  PVHLNRQPHAL-MMDDDQAANAATDSGATDDNSVLLPPPEETDSEEKPNKDGGDDGDNGV 118
KAF0754846.1_Ac        60  PVHLNRQPHAL-MMDDDQAANAATDSGATDDNSVLLPPPDETDGEEKPNKDGGDEEVNGG 118
XP_026816157.1_Rm      60  PVHLNRQPHGAMMMDDDQAATGAADSGATDDSSVLLPPPDETDNEEKPN-KDNADDEDNG 118
XP_025196103.1_Ms      60  PIHLGRHPHGAMMMDDDQAATAAADGGATDDSSVLLPPPDETDGEEKPNNKDNADGEDNG 119
XP_001949276.1_Ap      60  PVHLNRQQHGAMMMDDDQAANTAADSGATDDSSVLLPPPDDTENEEKPNKDNADDEDNGG 119
XP_022171233.1_Mp      60  PVHLNRQQHGAMMMDDDQAANTVAEGGATDDSSVLLPPPDETENEEKPNKDNADDEDNGG 119
XP_015366894.1_Dn      60  PVHLNRQQHGALMMDDDQAANTAADSGATDDSSVLLPPPDETDNEEKSNKDNADDEDNGG 119

Nemy_A_Dm              1   -------------------------------------------------MDTNSVSPLPP  11
VVC36571.1_Cc          116 NDAQWSEKPLGDDDDDDDQVPVLAMSGILPATATGGGGGGGGGGAGTGDSKKNVTGSNDP 175
KAE9533855.1_Agly      118 -----GDNNENRTGEDDDQVPVLAMSGILPATATGGGP--------ISDSKKNAGGPNDS 164
XP_027844671.1_Ago     119 -----EDNNENRAGEDDDQVPVLAMSGILPATATGGGP--------ISDSKKNAGGPNDS 165
KAF0754846.1_Ac        119 -----GDNNDNGAGEDDDQVPVLAMSGILPATATGGGP--------ISDSKKNAGGPNES 165
XP_026816157.1_Rm      119 -----GDNNDNRTGEDDDQVPVLAMSGILPATASGGGS--------ISDSKKNAGGPNDS 165
XP_025196103.1_Ms      120 -----GDNDDNGAAGDDDQVPVLAMSGILPATATGGGP--------ISDSKKNAGGPNDS 166
XP_001949276.1_Ap      120 -----DNN--DEAADDDDQVPVLAMSGILPATATGGGQ--------ISDSKKNAGGPNDS 164
XP_022171233.1_Mp      120 -----DNN--GEAADDDDQVPVLAMSGILPATATGGGP--------ISDSKKTAGGPNDS 164
XP_015366894.1_Dn      120 -----DNN--DEAADDDDQVPVLAMSGILPATATGGGP--------ISDSKKTSGGPNDT 164

Nemy_A_Dm              12  M---------EEAMDRVSEKSPPGTPNGIEMPPPPDEKRYEEDPDNAWNCCSWCEYLLIV  62
VVC36571.1_Cc          176 TSGGAGGTVTGNGENKSNEKSRYASGNGAGG-GTDQDDSGANNSQAPLGCGTTLEYVLLI 234
KAE9533855.1_Agly      165 LTGGAPG--TGNGENKSKEKSRYTTGSGTGA-GTDVEDSGTNNPQAALGCGTTFEYVVLV 221
XP_027844671.1_Ago     166 LTGGAPG--TGNGESKSKEKSRYTTGSGTGA-GTDVEDSGTNNPQAALGCGTTFEYVVLV 222
KAF0754846.1_Ac        166 LTGGAPG--TGNGENKSKEKSRYTAGSGTGA-GTDVEDSGTNNPQAALGCGTTFEYVVLV 222
XP_026816157.1_Rm      166 LTGGGTG--TGNGENKSKEKSRYTTGSGTGA-GTDVEDSGTNSPQAALGCGTTFEYVVLV 222
XP_025196103.1_Ms      167 LTGGGTGTGTGNGENKSKEKSRYTAGSGTGA-GTDVEDSGTSNPQAALGCGTTFEYVVLI 225
XP_001949276.1_Ap      165 A-GGGAG--TGNGENKSKEKSRYTTGSGTGA-GTDAEDSGTNNPQTALGCGTTFEYVVLV 220
XP_022171233.1_Mp      165 LTGGGTG--TGNGENKSKEKTRYTTGSGTGT-GTDAEDSGTNNPQAALGCGTTFEYVVLV 221
XP_015366894.1_Dn      165 LTGGGAG--TGNGENKSKEKTRYTAGSGTGT-GTDAEDSGTSNPQAALGCGTTFEYVVLV 221

                           Predicted Helix 1
Nemy_A_Dm              63  ILSTILLVGVLVLTLFWVMFYRDGFA-----WSE--------------------------  91
VVC36571.1_Cc          235 TTCTILLLSVTGTTIYWTMVYRGGYDCAWFPWPPQTPKQLLSVGIFTYKHHSQNSQNFSD 294
KAE9533855.1_Agly      222 TACSILLLAVTGTTVYWTMAYRGGYDPNWFPWPPQTPKQMLSVGVFTYKHP-QNFTA--- 277
XP_027844671.1_Ago     223 TACSILLLAVTGTTVYWTMAYRGGYDPNWFPWPPQTPKQMLSVGVFTYKHP-QNFTA--- 278
KAF0754846.1_Ac        223 TACSILLLAVTGTTVYWTMAYRGGYDPNWFPWPPQTPKQMLSVGVFTYKHP-QNFTT--- 278
XP_026816157.1_Rm      223 TACSILLLAVTGTTVYWTMAYRGGYDPNWFPWPPQTPKQMLSVGVFTYKHP-QNFTA--- 278
XP_025196103.1_Ms      226 TACSILLLAVTGTTVYWTMAYRGGYDPNWFPWPPQTPKQMLSVGVFTYKHP-QNFTT--- 281
XP_001949276.1_Ap      221 TACSFLLVAVTGTTVYWTMAYRGGYDPNWFPWPPQTPLQMLTIGVFTYKHP-QNITD--- 276
XP_022171233.1_Mp      222 TACSILLLAVTGTTVYWTMAYRGGYDTNWFPWPPLTPKQMLSGGVFTYKHPPQNFTT--- 278
XP_015366894.1_Dn      222 TACSILLLAVTGTTIYWTMAYRGGYDTNWFPWPPLTPREMLSGGVFTYKHP-QNFTT--- 277

                                                                     Predicted Helix 2
Nemy_A_Dm              92  ----------------------------------------------SPKQQFNLHPILMI 105
VVC36571.1_Cc          295 TNVDFGAGNSFGESFDGNYSTTSTIATSTPEQLSTQYTNKSSKIDLTDEQRFNLHPTLMT 354
KAE9533855.1_Agly      278 GGSDI--------------STTTIATTTVEQQQQSSQLNTSTSNVMIEDRRFNLHPTLMT 323
XP_027844671.1_Ago     279 GGSDI--------------STTTIATTTVEQQQQSSQLNTSTSNVLIEDRRFNLHPTLMT 324
KAF0754846.1_Ac        279 GSSDI--------------STTTIATTTVEQQQQSSQLNISTSNVLIEDRRFNLHPTLMT 324
XP_026816157.1_Rm      279 SGSDI--------------STTTIATTTTTEQQQSSQLNISSLNILIEDRRFNLHPTLMT 324
XP_025196103.1_Ms      282 SGSDI--------------STTTVTT-TTVEQQQSSQLNISSSNSLMEDRRFNLHPTLMT 326
XP_001949276.1_Ap      277 GGSDI--------------STTIS--TTTVEQQQSSLLNISSSNVSIEDQRFNLHPTLMT 320
XP_022171233.1_Mp      279 SASDI--------------STTVA--TTTVEQQQSSQLNISSSNFSIEDRRFNLHPTLMT 322
XP_015366894.1_Dn      278 GGSDI--------------STTIT--TTTVEQQQSSQLNISSSNVSIEDRRFNLHPTLMT 321
                                                                               *
```

### S4 Fig. continued

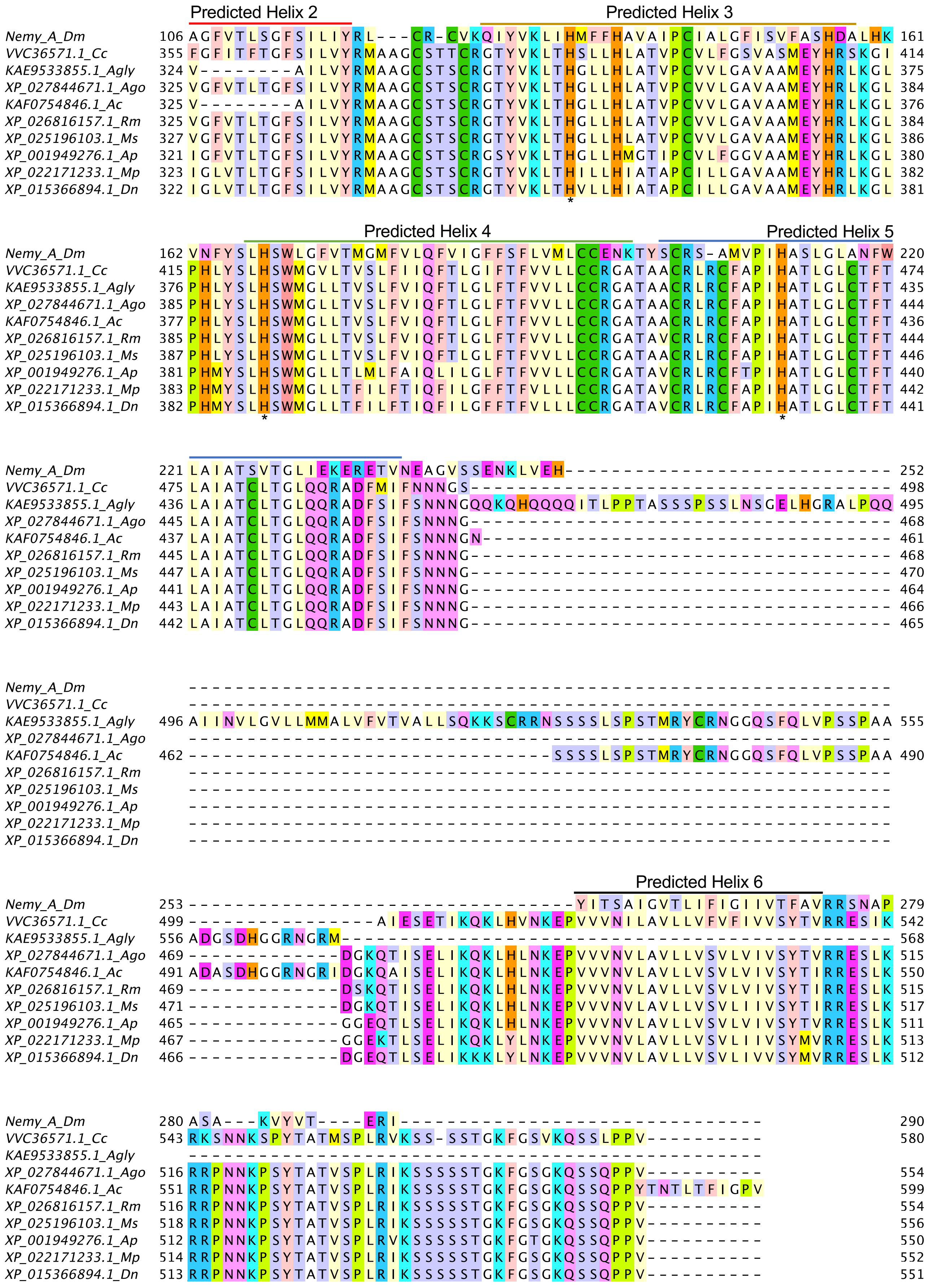
