## Supplemental Tables 1-8 for "Comparison of insect and human cytochrome b561 proteins: Insights into candidate ferric reductases in insects"

### S1 Table. Insect and outgroup cytb561 sequences used for this study.

| **Order** ***Species*** **(common name)** | **Accession number** | **Source/ "Query"** | **Common homologous sequence**[1] | **Group**[2] |
|---|---|---|---|---|
| Diptera *Drosophila melanogaster* (fruit fly) | NP_728727.1 (CG1275 isoform A) | FlyBase/ "Cytochrome b561" | G141-T276 | CG1275 |
| | NP_725208.1 (Nemy isoform A) | FlyBase/ "Cytochrome b561" | K94-I231 | Nemy |
| | NP_611079.2 (CG8399 isoform A) | FlyBase/ "Cytochrome b561" | R403-K536 | CG8399 |
| | NP_609982.1 (CG10165) | FlyBase/ "Cytochrome b561" | K43-Y177 | Group 4B |
| | NP_609990.1 (CG13077 isoform A) | FlyBase/ "Cytochrome b561" | S90-Q223 | Group 4B |
| | NP_609989.1 (CG13078) | FlyBase/ "Cytochrome b561" | S44-T178 | Group 4B |
| | NP_609986.1 (CG10337) | FlyBase/ "Cytochrome b561" | V43-N180 | Group 4B |
| | NP_570039.1 (CG3592) | FlyBase/ "Cytochrome b561" | H60-N189 | Group 4B |
| Diptera *Anopheles gambiae* (mosquito) | XP_315519.4 | NCBI BLAST/ "NP_728727.1" | S50-L186 | CG1275 |
| | XP_314126.2 | NCBI BLAST/ "NP_725208.1" | K63-T200 | Nemy |
| | XP_314065.2 | NCBI BLAST/ "NP_611079.2" | S22-G154 | CG8399 |
| | XP_314066.4 | NCBI BLAST/ "NP_611079.2" | K383-Q518 | CG8399 |
| | XP_320673.4 | NCBI BLAST/ "NP_728727.1" | M55-Y193 | Group 4A |
| | XP_001238089.2 | NCBI BLAST/ "NP_609982.1" | E54-E192 | Group 4B |
| | XP_552919.3 | NCBI BLAST/ "NP_609990.1" | G47-S185 | Group 4B |
| Hemiptera *Acyrthosiphon pisum* (aphid) | NP_001155374.1 | NCBI BLAST/ "NP_728727.1" | K54-S189 | CG1275 |
| | NP_001280323.1 | NCBI BLAST/ "NP_728727.1" | K57-C195 | CG1275 |
| | XP_001950854.1 | NCBI BLAST/ "NP_728727.1" | K53-C188 | CG1275 |
| | XP_003246890.1 | NCBI BLAST/ "NP_728727.1" | I76-S211 | CG1275 |
| | XP_001949276.1 | NCBI BLAST/ "NP_725208.1" | D309-Q451 | Nemy |
| | XP_001950579.2 | NCBI BLAST/ "NP_611079.2" | K554-T690 | CG8399 |
| Hymenoptera *Apis mellifera* (bee) | XP_006572086.1 | NCBI BLAST/ "NP_728727.1" | E51-N186 | CG1275 |
| | XP_396579.3 | NCBI BLAST/ "NP_611079.2" | N376-R510 | CG8399 |
| | XP_003249671.1 | NCBI BLAST/ "NP_609990.1" | N73-T203 | Group 4B |

| **Order**<br>***Species***<br>**(common name)** | **Accession number** | **Source/ "Query"** | **Common homologous sequence**[1] | **Group**[2] |
|---|---|---|---|---|
| | XP_001122176.1 | NCBI CDD/<br>"A. mellifera" +<br>"cytochrome_b_N" | I41-Y179 | Group 4A |
| Siphonaptera<br>*Ctenocephalides felis*<br>(flea) | XP_026462102.1 | NCBI BLAST/<br>"NP_728727.1" | E47-S187 | CG1275 |
| | XP_026473332.1 | NCBI BLAST/<br>"NP_725208.1" | K64-T203 | Nemy |
| | XP_026481553.1 | NCBI BLAST/<br>"NP_611079.2" | K382-K517 | CG8399 |
| | XP_026469480.1 | NCBI BLAST/<br>"NP_609990.1" | R68-F205 | Group 4A |
| | XP_026469242.1 | NCBI CDD/<br>"C. felis" +<br>"cyt_b561" | I68-F206 | Group 4A |
| Lepidoptera<br>*Papilio xuthus*<br>(butterfly) | NP_001298968.1 | NCBI BLAST/<br>"NP_725208.1" | E66-T203 | Nemy |
| | XP_013164083.1 | NCBI BLAST/<br>"NP_611079.2" | K390-Y526 | CG8399 |
| | XP_013172799.1 | NCBI BLAST/<br>"NP_609982.1" | G70-D203 | Group 4B |
| | XP_013172813.1 | NCBI BLAST/<br>"NP_609982.1" | G69-D201 | Group 4B |
| | XP_013162691.1 | NCBI BLAST/<br>"NP_609990.1" | R56-H188 | Group 4B |
| | XP_013174961.1 | NCBI CDD/<br>"P. xuthus" +<br>"cyt_b561" | T64-L201 | Group 4A |
| | XP_013162686.1 | NCBI CDD/<br>"P. xuthus" +<br>"cytochrome_b_N" | I87-N221 | Group 4B |
| | XP_013172802.1 | NCBI CDD/<br>"P. xuthus" +<br>"cytochrome_b_N" | N74-Y202 | Group 4B |
| | XP_013172804.1 | NCBI CDD/<br>"P. xuthus" +<br>"cytochrome_b_N" | I60-Y192 | Group 4B |
| Psocodea<br>*Pediculus humanus corporis*<br>(louse) | XP_002426701.1 | NCBI BLAST/<br>"NP_728727.1" | K45-T181 | CG1275 |
| | XP_002430226.1 | NCBI BLAST/<br>"NP_725208.1" | K88-T223 | Nemy |
| | XP_002423127.1 | NCBI BLAST/<br>"NP_611079.2" | D396-K532 | CG8399 |
| Coleoptera<br>*Tribolium castaneum*<br>(beetle) | XP_008194670.1 | NCBI BLAST/<br>"NP_728727.1" | Q60-T195 | CG1275 |
| | XP_008198104.1 | NCBI BLAST/<br>"NP_725208.1" | I106-T245 | Nemy |
| | XP_015836986.1 | NCBI BLAST/<br>"NP_611079.2" | K371-K506 | CG8399 |
| | XP_015837014.1 | NCBI BLAST/<br>"NP_609982.1" | D43-E178 | Group 4B |
| | XP_008201603.1 | NCBI BLAST/<br>"NP_609990.1" | Q113-Y250 | Group 4A |

| **Order**<br>***Species***<br>**(common name)** | **Accession number** | **Source/**<br>**“Query”** | **Common homologous sequence**[1] | **Group**[2] |
|---|---|---|---|---|
| | XP_974632.1 | NCBI CDD/<br>“T. castaneum” +<br>“cyt_b561” | N49-D184 | Group 4B |
| | XP_974652.1 | NCBI CDD/<br>“T. castaneum" +<br>“cytochrome_b_N” | D42-L176 | Group 4B |
| | XP_008195477.1<br>(edited) | NCBI CDD/<br>“T. castaneum" +<br>“cytochrome_b_N” | Y39-Y175 | Group 4B |
| Blattodea<br>*Zootermopsis nevadensis*<br>(termite) | XP_021935166.1 | NCBI BLAST/<br>“NP_728727.1” | D47-T182 | CG1275 |
| | XP_021939496.1 | NCBI BLAST/<br>“NP_725208.1” | K118-T258 | Nemy |
| | XP_021919699.1 | NCBI BLAST/<br>“NP_611079.2” | N377-K512 | CG8399 |
| | XP_021927739.1 | NCBI CDD/<br>“Z. nevadensis” +<br>“cyt_b561” | Y73-Y211 | Group 4A |
| Phylum Placozoa \| *Trichoplax adhaerens* | XP_002110723.1 | NCBI BLAST/<br>“NP_728727.1” | F56-F191 | Outgroup |

[1]The common homologous sequence is the amino acid sequence that corresponds to human Dcytb A44-T179. This region includes most of the core cytb561 domain of the protein.
[2]Sequences were assigned to a group based on phylogenetic and sequence analyses. The *D. melanogaster* protein name was used for all insect orthologs that group with CG1275, Nemy, or CG8399. Seven Group 4 sequences are more similar to human TScytb than any *D. melanogaster* cytb561s and are referred to as subgroup 4A, whereas the remaining Group 4 proteins belong to subgroup 4B. (Note that the 4A and 4B subgroups are based on sequence comparisons rather than phylogenetic relationship.)

### S2 Table. Sequence identity between insect and *D. melanogaster* cytb561s.

| | | | *D. melanogaster* Sequences | | | | | | | |
|---|---|---|---|---|---|---|---|---|---|---|
| **Species** | **Group** | **Accession Number** | **CG1275** | **Nemy** | **CG8399** | **CG10165** | **CG13077** | **CG13078** | **CG10337** | **CG3592** |
| *Dm* | CG1275 | NP_728727.1 | **100**[1] | 39.3 | 18.9 | 17 | 16.3 | 16.9 | 20.3 | 19.6 |
| *Ag* | CG1275 | XP_315519.4 | **68.6** | 35.5 | 17.4 | 16.9 | 16.2 | 16.1 | 16 | 17.9 |
| *Ap* | CG1275 | NP_001155374.1 | **54.4** | 42.1 | 14.7 | 21.3 | 17 | 15.5 | 16.8 | 18 |
| *Ap* | CG1275 | NP_001280323.1 | **43.6** | 31.9 | 15 | 16.6 | 14.5 | 13.7 | 21.7 | 15.5 |
| *Ap* | CG1275 | XP_001950854.1 | **48.5** | 33.6 | 19.6 | 15.6 | 16.3 | 17.6 | 16.8 | 17.4 |
| *Ap* | CG1275 | XP_003246890.1 | **48.5** | 33.6 | 16 | 17 | 15.6 | 18.3 | 18.1 | 16.5 |
| *Am* | CG1275 | XP_006572086.1 | **54.4** | 36.4 | 18.2 | 17 | 19.1 | 19.7 | 19.4 | 20.1 |
| *Cf* | CG1275 | XP_026462102.1 | **51.8** | 30.3 | 17.6 | 17.8 | 16.4 | 16.3 | 18.2 | 16.8 |
| *Ph* | CG1275 | XP_002426701.1 | **58.4** | 36.2 | 16.4 | 16 | 16 | 18.6 | 17.5 | 19.9 |
| *Tc* | CG1275 | XP_008194670.1 | **56.6** | 38.6 | 18.2 | 19.1 | 14.9 | 17.7 | 16.8 | 20.3 |
| *Zn* | CG1275 | XP_021935166.1 | **58.8** | 34.3 | 18.9 | 21.3 | 17 | 18.3 | 16.1 | 21 |
| *Dm* | Nemy | NP_725208.1 | 39.3 | **100** | 13.1 | 17 | 19.1 | 18.3 | 14.3 | 17.6 |
| *Ag* | Nemy | XP_314126.2 | 42.1 | **67.4** | 19.3 | 16.3 | 15.6 | 16.9 | 13.6 | 18.3 |
| *Ap* | Nemy | XP_001949276.1 | 36.6 | **46.9** | 22.6 | 14.8 | 19 | 20.3 | 17.4 | 22.2 |
| *Cf* | Nemy | XP_026473332.1 | 45 | **63.6** | 15.6 | 16.8 | 18.2 | 22.9 | 17.7 | 20.4 |
| *Px* | Nemy | NP_001298968.1 | 42.1 | **54.3** | 14.5 | 17.7 | 18.4 | 20.4 | 17 | 19 |
| *Ph* | Nemy | XP_002430226.1 | 41.4 | **52.9** | 16.3 | 14.7 | 16.8 | 19.4 | 15 | 14.8 |
| *Tc* | Nemy | XP_008198104.1 | 45.7 | **61.4** | 17 | 18.2 | 16.1 | 20.1 | 18.4 | 20.4 |
| *Zn* | Nemy | XP_021939496.1 | 42.6 | **62.4** | 16.7 | 15.1 | 15.8 | 19 | 16.3 | 19.3 |
| *Dm* | CG8399 | NP_611079.2 | 18.9 | 13.1 | **100** | 12.1 | 12.1 | 13.4 | 11.6 | 11.3 |
| *Ag* | CG8399 | XP_314065.2 | 18.1 | 13.7 | **40.3** | 14.1 | 14.1 | 16.9 | 13.5 | 12.6 |
| *Ag* | CG8399 | XP_314066.4 | 19.4 | 18.5 | **65.4** | 13.6 | 17.1 | 20.7 | 12.2 | 13.3 |
| *Ap* | CG8399 | XP_001950579.2 | 15.6 | 16.8 | **54.7** | 11.3 | 13.4 | 14.7 | 10.9 | 15.3 |
| *Am* | CG8399 | XP_396579.3 | 14.7 | 15.2 | **53.3** | 12.2 | 12.2 | 19.3 | 9.5 | 9.2 |
| *Cf* | CG8399 | XP_026481553.1 | 20.1 | 19.2 | **62.5** | 11.4 | 12.9 | 15.7 | 12.2 | 11.9 |
| *Px* | CG8399 | XP_013164083.1 | 16.8 | 15.9 | **62** | 14.9 | 12.1 | 13.4 | 15 | 9.9 |
| *Ph* | CG8399 | XP_002423127.1 | 18.9 | 16.7 | **62** | 9.7 | 11 | 13.7 | 10.9 | 10.2 |
| *Tc* | CG8399 | XP_015836986.1 | 16 | 17.1 | **65.4** | 12.1 | 12.1 | 16.4 | 10.8 | 10.5 |
| *Zn* | CG8399 | XP_021919699.1 | 19.4 | 16.4 | **64** | 10.7 | 12.1 | 15.7 | 10.8 | 9.8 |
| *Dm* | Group 4B | NP_609982.1 | 16.9 | 16.9 | 12 | **100** | 25.9 | 27.9 | 21.9 | 19.3 |
| *Dm* | Group 4B | NP_609990.1 | 16.2 | 19 | 12 | 25.9 | **100** | 51.1 | 26 | 20.7 |
| *Dm* | Group 4B | NP_609989.1 | 16.8 | 18.2 | 13.3 | 27.9 | 51.1 | **100** | 22.4 | 17 |
| *Dm* | Group 4B | NP_609986.1 | 20.1 | 14.2 | 11.5 | 21.9 | 26 | 22.4 | **100** | 34.1 |
| *Dm* | Group 4B | NP_570039.1 | 19.4 | 17.5 | 11.2 | 19.3 | 20.7 | 17 | 34.1 | **100** |
| *Ag* | Group 4A | XP_320673.4[1] | 22.9 | 19.4 | 21.5 | **27.3** | 20.3 | 20.3 | 17.8 | 15.8 |
| *Ag* | Group 4B | XP_001238089.2 | 19 | 16.1 | 18.9 | **30.2** | 27.5 | 23.9 | 18.5 | 18 |
| *Ag* | Group 4B | XP_552919.3 | 16.2 | 14.1 | 13.3 | 27.3 | **29** | 26.1 | 22.8 | 22.5 |
| *Am* | Group 4A | XP_001122176.1[2] | 19 | 16.7 | 14 | 20.1 | 19.4 | 19.4 | 17.1 | **21.6** |
| *Am* | Group 4B | XP_003249671.1 | 21.8 | 20.1 | 14 | 28.1 | 31.9 | **34.1** | 26.2 | 21 |
| *Cf* | Group 4A | XP_026469480.1[2] | 21.3 | 16.8 | 16.2 | **21.6** | 21 | 15.9 | 18.5 | 16.5 |
| *Cf* | Group 4A | XP_026469242.1[2] | **22** | 20.4 | 17.6 | 21.6 | 19.4 | 16.4 | 17.9 | 14.5 |
| *Px* | Group 4A | XP_013174961.1[2] | 21.8 | 21.5 | 17.5 | **24.5** | 19.6 | 23.2 | 15.8 | 14.4 |
| *Px* | Group 4B | XP_013172799.1 | 15.6 | 15.5 | 10.6 | **28.1** | 27.6 | 25.9 | 20.7 | 18 |
| *Px* | Group 4B | XP_013172813.1 | 17.9 | 17 | 10.6 | **28.1** | 26.1 | 26.7 | 18.6 | 16.5 |
| *Px* | Group 4B | XP_013162691.1 | 17.6 | 18.3 | 16.8 | 24.4 | **24.6** | 24.4 | 20.5 | 18.7 |
| *Px* | Group 4B | XP_013162686.1 | 13.4 | 12 | 17.6 | **24.4** | 23.1 | 22.2 | 17.8 | 17.1 |
| *Px* | Group 4B | XP_013172802.1 | 15 | 19.7 | 9.4 | 20.9 | 19.4 | **21.4** | 19.7 | 16.1 |
| *Px* | Group 4B | XP_013172804.1 | 16.9 | 18.3 | 16.9 | 24.4 | 25.4 | **27.4** | 21.2 | 16.4 |
| *Tc* | Group 4A | XP_008201603.1[2] | 21.3 | 19.6 | 12.7 | **25.9** | 20.9 | 19.4 | 19.3 | 17.4 |
| *Tc* | Group 4B | XP_008195477.1[3] | 17.5 | 16.9 | 18.1 | 22.6 | **27.9** | 24.1 | 20 | 17.9 |
| *Tc* | Group 4B | XP_015837014.1 | 20.4 | 16.2 | 16.8 | 24.3 | **30.4** | 22.2 | 22.8 | 17.9 |
| *Tc* | Group 4B | XP_974632.1 | 17.6 | 14.8 | 15.4 | 19.9 | **27.4** | 19.3 | 20.7 | 19.3 |
| *Tc* | Group 4B | XP_974652.1 | 17 | 19 | 14.8 | 26.5 | **30.4** | 27.4 | 22.8 | 21.7 |
| *Zn* | Group 4A | XP_021927739.1[2] | 19.9 | 21.1 | 16.2 | 25.2 | **30.2** | 18.6 | 20 | 17.4 |

[1]Sequence identity (as a percent) between the homologous region of each insect cytb561 and each *D. melanogaster* cytb561. The highest identity in each row is cell shaded and listed in bold. Group 4A sequences are shaded in pale orange and all other sequences are shaded in pale blue.
[2]Group 4A insect sequences with higher identity to a human cytb561 than a *D. melanogaster* cytb561.
[3]Edited to correct a gene prediction error.

**S3 Table. ChimeraX-predicted contacts for Dcytb, CG1275, and Nemy with heme and ascorbate molecules.**

| Ligand[1] | Position[2] | Dcytb Amino Acid | # Contacts[3] | # H-bonds[3] | Nemy Amino Acid | # Contacts | # H-bonds | CG1275 Amino Acid | # Contacts | # H-Bonds | Jalview Score[4] |
|---|---|---|---|---|---|---|---|---|---|---|---|
| Heme (NC) | H1-2 | Trp30 | 3 | | Trp79 | 1 | | Trp126 | 1 | | * |
| Heme (NC) | H1-3 | Phe47 | 5 | | Phe97 | 10 | | Phe144 | 12 | | * |
| Ascorbate (NC) | H1-3 | Phe47 | 2 | | Phe97 | 3 | | Phe144 | 2 | | * |
| Heme (NC) | H1 | His50 | 23 | 2 | His100 | 19 | 3 | His147 | 20 | 2 | * |
| Heme (NC) | H1+1 | Pro51 | 4 | | Pro101 | 4 | | Pro 148 | 6 | | * |
| Heme (NC) | H1+4 | Met54 | 8 | | Met104 | 7 | | Met 151 | 10 | | * |
| Heme (NC) | H1+8 | Phe58 | 1 | | Phe108 | 3 | | Phe155 | 3 | | 1 |
| Heme (C) | H1+12 | Gln62 | 0 | | Ser112 | 0 | | Tyr159 | 4 | | 4 |
| Heme (C) | H1+16 | Ile66 | 3 | | Ile116 | 2 | | Ile163 | 5 | | 8 |
| Heme (C) | H1+19 | Tyr69 | 14 | | Tyr119 | 9 | | Tyr166 | 7 | | * |
| Heme (C) | H1+20 | Arg70 | 5 | 1 | Arg120 | 0 | | Arg167 | 4 | 1 | 9 |
| Heme (C) | H2-7 | Lys79 | 6 | | Gln129 | 0 | | Lys174 | 0 | | 3 |
| Ascorbate (C) | H2-7 | Lys 79 | 6 | 1 | Gln129 | 0 | | Lys174 | 4 | 1 | 3 |
| Heme (C) | H2-3 | Lys83 | 4 | | Lys131 | 3 | | Lys178 | 1 | | * |
| Ascorbate (C) | H2-3 | Lys 83 | 4 | 2 | Lys131 | 6 | 2 | Lys178 | 4 | 2 | * |
| Heme (C) | H2 | His86 | 23 | 1 | His134 | 15 | 1 | His181 | 21 | 2 | * |
| Heme (C) | H2+1 | Ala87 | 2 | | Met135 | 1 | | Ala182 | 1 | | 5 |
| Heme (C) | H2+4 | Asn90 | 7 | | His138 | 12 | | His185 | 17 | | 4 |
| Heme (NC) | H2+15 | Val101 | 1 | | Phe149 | 2 | | Leu 196 | 3 | | 6 |
| Heme (NC) | H2+18 | Val104 | 5 | | Val152 | 8 | | Val199 | 10 | | 8 |
| Heme (NC) | H2+19 | Phe105 | 2 | | Phe153 | 8 | | Phe200 | 6 | | 7 |
| Heme (NC) | H2+22 | His108 | 1 | | His156 | 8 | | His203 | 7 | | * |
| Ascorbate (NC) | H2+22 | His108 | 2 | 1 | His156 | 2 | 1 | His203 | 1 | 1 | * |
| Ascorbate (NC) | H3-7 | Ile113 | 0 | | Lys161 | 23 | 1 | Ile210 | 0 | | 6 |
| Heme (NC) | H3-5 | Asn115 | 4 | | Asn163 | 5 | | Asn212 | 5 | | 5 |
| Heme (NC) | H3-4 | Met116 | 1 | 1 | Phe164 | 0 | 1 | Met213 | 1 | 1 | 8 |
| Heme (NC) | H3-3 | Tyr117 | 1 | | Tyr165 | 0 | | Tyr214 | 0 | | * |
| Heme (NC) | H3-2 | Ser118 | 4 | 2 | Ser166 | 4 | 1 | Ser215 | 4 | 1 | 8 |
| Heme (NC) | H3 | His120 | 16 | | His168 | 30 | | His217 | 24 | | * |
| Heme (NC) | H3+1 | Ser121 | 3 | | Ser169 | 9 | | Ser218 | 1 | | * |
| Heme (NC) | H3+4 | Gly124 | 4 | | Gly172 | 6 | | Gly221 | 4 | | * |
| Heme (NC) | H3+5 | Leu125 | 0 | | Phe173 | 1 | | Leu222 | 1 | | 7 |
| Heme (NC) | H3+7 | Ala127 | 0 | | Thr175 | 1 | | Ala224 | 0 | | 7 |

| Ligand[1] | Position[2] | Dcytb Amino Acid | # Contacts[3] | # H-bonds[3] | Nemy Amino Acid | # Contacts | # H-bonds | CG1275 Amino Acid | # Contacts | # H-Bonds | Jalview Score[4] |
|---|---|---|---|---|---|---|---|---|---|---|---|
| Heme (NC) | H3+8 | Val128 | 1 | | Met176 | 0 | | Val225 | 1 | | 8 |
| Heme (C) | H3+11 | Tyr131 | 8 | | Phe179 | 3 | | Phe228 | 6 | | 9 |
| Heme (C) | H3+14 | Gln134 | 6 | | Gln182 | 2 | | Gln231 | 0 | | * |
| Heme (C) | H3+15 | Leu135 | 1 | | Phe183 | 4 | | Tyr232 | 4 | | 7 |
| Heme (C) | H3+18 | Gly138 | 4 | | Gly186 | 10 | | Gly235 | 5 | | * |
| Heme (C) | H3+19 | Phe139 | 3 | | Phe187 | 4 | | Phe236 | 4 | | 6 |
| Heme (C) | H3+22 | Phe142 | 5 | | Phe190 | 13 | | Phe239 | 7 | | 9 |
| Ascorbate (C) | H3+22 | Phe142 | 4 | | Phe190 | 6 | | Phe239 | 5 | | 9 |
| Heme (C) | H3+23 | Leu143 | 5 | | Leu191 | 0 | | Leu240 | 4 | | 6 |
| Ascorbate (C) | H4-7 | Arg152 | 5 | 1 | Arg204 | 10 | 3 | Arg249 | 6 | | 9 |
| Ascorbate (C) | H4-6 | Ala153 | 0 | | Ser205 | 9 | 1 | Ile250 | 4 | | 3 |
| Heme (C) | H4-3 | Met156 | 3 | | Val208 | 0 | | Met253 | 3 | | 5 |
| Ascorbate (C) | H4-3 | Met156 | 1 | | Val208 | 6 | | Met253 | 8 | | 5 |
| Heme (C) | H4 | His159 | 14 | | His211 | 31 | 2 | His256 | 20 | 1 | * |
| Heme (C) | H4+1 | Val160 | 2 | | Ala212 | 4 | | Ile257 | 2 | | 5 |
| Heme (C) | H4+4 | Gly163 | 1 | | Gly215 | 2 | | Gly260 | 3 | | * |
| Heme (C) | H4+5 | Ile164 | 1 | | Leu216 | 0 | | Leu261 | 2 | | 9 |
| Heme (NC) | H4+11 | Val170 | 2 | | Ala222 | 1 | | Ala267 | 1 | | 8 |
| Heme (NC) | H4+14 | Thr173 | 4 | | Thr225 | 12 | | Ser270 | 2 | | 7 |
| Heme (NC) | H4+15 | Ala174 | 3 | | Ser226 | 7 | | Ala271 | 3 | | 7 |
| Heme (NC) | H4+18 | Gly177 | 5 | | Gly229 | 5 | | Gly274 | 11 | | * |
| Heme (NC) | H4+19 | Leu178 | 2 | | Leu230 | 1 | | Ile275 | 6 | | 8 |
| Heme (NC) | H4+21 | Glu180 | 7 | | Glu232 | 9 | | Glu277 | 7 | | 8 |
| Heme (NC) | H4+22 | Lys181 | 2 | | Lys233 | 2 | | Lys278 | 2 | | 9 |
| Ascorbate (NC) | H4+25 | Phe184 | 4 | | Glu236 | 29 | | Phe281 | 2 | | 4 |
| Heme (C) | n/a | Val219 | 2 | | Val273 | 1 | | Ala316 | 0 | | 7 |
| Heme (C) | n/a | Lys225 | 5 | 2 | Pro279 | 0 | | Lys332 | 0 | | 3 |

[1]Ligand: (C) = cytoplasmic; (NC) = non-cytoplasmic.
[2]Position: Relative to H1-H4 (heme-coordinating histidines).
[3]Number of contacts and H-bonds separately predicted using ChimeraX.
[4]Jalview Scores are based on the Figure 4 alignment. Higher scores correspond to higher conservation with an * for entirely conserved.

**S4 Table. AlphaFold model confidence for *D. melanogaster* CG1275, Nemy, and CG8399 structures based on per-residue confidence score (pLDDT) between 0 and 100.**

| Region of<br>CG1275 Isoform A | % Very Low<br>(pLDDT <50) | % Low<br>(70 > pLDDT > 50) | % Confident<br>(90 > pLDDT > 70) | % Very High<br>(pLDDT > 90) |
|---|---|---|---|---|
| N-terminus<br>Amino acids 1-99 | 93.9% (93/99) | 5.1% (5/99) | 1.0% (1/99) | 0% (0/99) |
| Full cytb561 domain<br>Amino acids 100-317 | 0% (0/218) | 0% (0/218) | 2.3% (5/218) | 97.7% (213/218) |
| C-terminus<br>Amino acids 318-340 | 4.4% (1/23) | 39.1% (9/23) | 30.4% (7/23) | 26.1% (6/23) |

| Region of<br>Nemy Isoform A | % Very Low<br>(pLDDT <50) | % Low<br>(70 > pLDDT > 50) | % Confident<br>(90 > pLDDT > 70) | % Very High<br>(pLDDT > 90) |
|---|---|---|---|---|
| N-terminus<br>Amino acids 1-53 | 75.5% (40/53) | 18.9% (10/53) | 5.7% (3/53) | 0% (0/53) |
| Full cytb561 domain<br>Amino acids 54-274 | 0% (0/221) | 0% (0/221) | 17.2% (38/221) | 82.8% (183/221) |
| C-terminus<br>Amino acids 275-290 | 87.5% (14/16) | 6.25% (1/16) | 6.25% (1/16) | 0% (0/16) |

| Region of<br>CG8399 Isoform A | % Very Low<br>(pLDDT <50) | % Low<br>(70 > pLDDT > 50) | % Confident<br>(90 > pLDDT > 70) | % Very High<br>(pLDDT > 90) |
|---|---|---|---|---|
| Signal peptide<br>Amino acids 1-30 | 76.7% (23/30) | 23.3% (7/30) | 0% (0/30) | 0% (0/30) |
| Reeler domain<br>Amino acids 31-184 | 0% (0/154) | 2.6% (4/154) | 40.3% (62/154) | 57.1% (88/154) |
| Linker (between reeler and DOMON domains)<br>Amino acids 185-223 | 87.2% (34/39) | 10.3% (4/39) | 2.6% (1/39) | 0% (0/39) |
| DOMON domain<br>Amino acids 224-400 | 0% (0/177) | 0% (0/177) | 37.9% (67/177) | 62.1% (110/177) |
| Linker (between DOMON & cytb561 domains)<br>Amino acids 401-403 | 0% (0/3) | 0% (0/3) | 100% (3/3) | 0% (0/3) |
| Full cytb561 domain<br>Amino acids 404-636 | 0% (0/233) | 9.87% (23/233) | 21.46% (50/233) | 68.67% (160/233) |
| Cytb561 core domain[1] (helices 1-4 and their loops)<br>Amino acids 404-535 | 0% (0/132) | 0% (0/132) | 18.2% (24/132) | 81.8% (108/132) |
| C-terminus<br>Amino acids 637-647 | 27.3% (3/11) | 36.4% (4/11) | 36.4% (4/11) | 0% (0/11) |

[1]A subset of the residues in the row above to emphasize that the core domain prediction is much better than the full cytb561 domain prediction.

**S5 Table. Insect sequences with similar extended regions to *A. pisum* Nemy.**

| **Order**<br>***Species* (common name)** | **Accession number[1]** |
|---|---|
| Hemiptera<br>*Aphis craccivora* (cowpea aphid) | KAF0754846.1 |
| Hemiptera<br>*Aphis glycines* (soybean aphid) | KAE9533855.1 |
| Hemiptera<br>*Aphis gossypii* (cotton aphid) | XP_027844671.1 |
| Hemiptera<br>*Cinara cedri* (cedar bark aphid) | VVC36571.1 |
| Hemiptera<br>*Diuraphis noxia* (Russian wheat aphid) | XP_015366894.1 |
| Hemiptera<br>*Melanaphis sacchari* (sugarcane aphid) | XP_025196103.1 |
| Hemiptera<br>*Myzus persicae* (green peach aphid) | XP_022171233.1 |
| Hemiptera<br>*Rhopalosiphum maidis* (corn aphid) | XP_026816157.1 |
| Hemiptera<br>*Sipha flava* (yellow sugarcane aphid) | XP_025407957.1 |

[1]Sequences identified from a BLAST search using query "XP_00194292276.1," the *A. pisum* Nemy sequence with approximately 150 additional N-terminal amino acids and an extended non-cytoplasmic loop connecting helix 1 and helix 2, against the class Insecta.

**S6 Table. Insect sequences with two DOMON domains.**

| Order<br>*Species* (common name) | Accession number[1] |
|---|---|
| Hemiptera<br>*Aphis craccivora* (cowpea aphid) | KAF0772237.1 |
| Hemiptera<br>*Aphis glycines* (soybean aphid) | KAE9523894.1 |
| Hemiptera<br>*Aphis gossypii* (melon aphid) | XP_027841628.1 |
| Hemiptera<br>*Cinara cedri* (cedar bark aphid) | VVC24734.1 |
| Hemiptera<br>*Diuraphis noxia* (Russian wheat aphid) | XP_015375719.1 |
| Hemiptera<br>*Melanaphis sacchari* (sugarcane aphid) | XP_025198216.1 |
| Hemiptera<br>*Rhopalosiphum maidis* (corn aphid) | XP_026806269.1 |
| Hemiptera<br>*Sipha flava* (yellow sugarcane aphid) | XP_025420155.1 |
| Hemiptera<br>*Myzus persicae* (green peach aphid) | XP_022176482.1 |
| Hemiptera<br>*Apolygus lucorum* (small green plant bug) | KAF6216808.1 |
| Hemiptera<br>*Bemisia tabaci* (silverleaf whitefly) | XP_018904139.1 |
| Hemiptera<br>*Cimex lectularius* (bed bug) | XP_014245063.1 |
| Hemiptera<br>*Diaphorina citri* (Asian citrus psyllid) | XP_017300331.1 |
| Hemiptera<br>*Laodelphax striatellus* (small brown planthopper) | RZF35324.1 |
| Hemiptera<br>*Halyomorpha halys* (brown marmorated stink bug) | XP_014284211.1 |
| Hemiptera<br>*Nilaparvata lugens* (brown planthopper) | XP_039277083.1 |
| Hemiptera<br>*Homalodisca vitripennis* (glassy-winged sharpshooter) | KAG8312720.1 |
| Coleoptera<br>*Diabrotica vergifera* (corn rootworm) | XP_028141846.1 |
| Coleoptera<br>*Leptinotarsa decemlineata* (Colorado potato beetle) | XP_023022567.1 |

[1]Sequences identified from a BLAST search using query "XP_001950579.2," the *A. pisum* CG8399 sequence with two DOMON domains, against the class Insecta.

**S7 Table. Single-domain insect sequences with highest similarity to CG8399.**

| **Order**<br>***Species* (common name)** | **Accession number[1]** |
|---|---|
| Diptera<br>*Aedes aegypti* (Yellow fever mosquito) | XP_021698251.1 |
| Diptera<br>*Aedes albopictus* (Asian tiger mosquito) | XP_029708302.1 |
| Diptera<br>*Anopheles albimanus* (New world malaria mosquito) | XP_035793296.1 |
| Diptera<br>*Anopheles arabiensis* (African malaria mosquito) | XP_040158133.1 |
| Diptera<br>*Anopheles coluzzii* (African malaria mosquito) | XP_040226982.1 |
| Diptera<br>*Anopheles darlingi* (American malaria mosquito) | ETN60734.1 |
| Diptera<br>*Anopheles merus* (mosquito) | XP_041769842.1 |
| Diptera<br>*Anopheles sinensis* (mosquito) | KFB44363.1 |
| Diptera<br>*Anopheles stephensi* (Indo-Pakistan malaria mosquito) | XP_035914634.1 |
| Diptera<br>*Culex quinquefasciatus* (southern house mosquito) | XP_038118957.1 |
| Diptera<br>*Bradysia coprophila* (darkwinged fungus gnat) | XP_037036556.1 |
| Diptera<br>*Bradysia odoriphaga* (darkwinged fungus gnat) | KAG4076368.1 |

[1]Sequences identified from a BLAST search using query "XP_314065.2," the *A. gambiae* CG8399 single-domain cytb561 sequence lacking both reeler and DOMON domains, against the class Insecta.

**S8 Table. Predicted subcellular localization of insect cytb561 sequences based on DEEPLOC-1.0 webserver.**

| Accession number | Group | Predicted Localization | Cell membrane[1] | ER[2] | Golgi | Lysosome |
|---|---|---|---|---|---|---|
| XP_015837014.1_Tc | Group 4 | ER | 0.1201 | 0.3888 | 0.2281 | 0.2626 |
| XP_974632.1_Tc | Group 4 | Golgi | 0.0647 | 0.2818 | 0.3967 | 0.2562 |
| XP_974652.1_Tc | Group 4 | ER | 0.0748 | 0.4115 | 0.2891 | 0.2244 |
| XP_008195477.1_Tc | Group 4 | Golgi | 0.0352 | 0.2676 | 0.5114 | 0.1855 |
| XP_552919.3_Ag | Group 4 | Golgi | 0.1246 | 0.202 | 0.3427 | 0.3298 |
| XP_001238089.2_Ag | Group 4 | Golgi | 0.1175 | 0.2737 | 0.3197 | 0.2855 |
| XP_013172802.1_Px | Group 4 | Lysosome | 0.0786 | 0.1172 | 0.2758 | 0.5216 |
| XP_013162686.1_Px | Group 4 | Lysosome | 0.1012 | 0.1563 | 0.3461 | 0.3942 |
| XP_013172799.1_Px | Group 4 | Golgi | 0.1573 | 0.1845 | 0.3784 | 0.2769 |
| XP_013172813.1_Px | Group 4 | Golgi | 0.106 | 0.2345 | 0.3583 | 0.2996 |
| XP_013162691.1_Px | Group 4 | Lysosome | 0.0933 | 0.3079 | 0.2671 | 0.3288 |
| XP_013172804.1_Px | Group 4 | Golgi | 0.0955 | 0.3178 | 0.3287 | 0.2489 |
| NP_609982.1_Dm | Group 4 | Golgi | 0.1152 | 0.2663 | 0.3303 | 0.2876 |
| NP_609986.1_Dm | Group 4 | Lysosome | 0.0881 | 0.2983 | 0.2308 | 0.3821 |
| NP_570039.1_Dm | Group 4 | Lysosome | 0.1743 | 0.1386 | 0.2253 | 0.4613 |
| XP_003249671.1_Am | Group 4 | Golgi | 0.143 | 0.1862 | 0.3732 | 0.2959 |
| NP_001260592.1_Dm | Group 4 | Golgi | 0.0811 | 0.1634 | 0.4081 | 0.3457 |
| NP_609989.1_Dm | Group 4 | Golgi | 0.0549 | 0.2851 | 0.3584 | 0.3006 |
| XP_001122176.1_Am | Group 4A | Lysosome | 0.1158 | 0.191 | 0.3412 | 0.3515 |
| XP_026469480.1_Cf | Group 4A | Lysosome | 0.0894 | 0.2834 | 0.2528 | 0.3723 |
| XP_026469242.1_Cf | Group 4A | Lysosome | 0.1185 | 0.1749 | 0.2805 | 0.423 |
| XP_008201603.1_Tc | Group 4A | ER | 0.099 | 0.3092 | 0.297 | 0.2901 |
| XP_320673.4_Ag | Group 4A | Golgi | 0.0653 | 0.2733 | 0.3963 | 0.2633 |
| XP_013174961.1_Px | Group 4A | Lysosome | 0.1475 | 0.171 | 0.2854 | 0.3939 |
| XP_021927739.1_Zn | Group 4A | Lysosome | 0.1565 | 0.1701 | 0.2632 | 0.4072 |
| XP_314065.2_Ag | CG8399 | Lysosome | 0.3707 | 0.04 | 0.0956 | 0.4929 |
| XP_001950579.2_Ap | CG8399 | Cell membrane | 0.8756 | 0.0243 | 0.0083 | 0.0854 |
| XP_396579.3_Am | CG8399 | Cell membrane | 0.8254 | 0.0605 | 0.0097 | 0.0967 |
| NP_611079.2_Dm | CG8399 | Cell membrane | 0.8186 | 0.0515 | 0.0132 | 0.1102 |
| XP_013164083.1_Px | CG8399 | Cell membrane | 0.7493 | 0.0731 | 0.0383 | 0.125 |
| XP_002423127.1_Ph | CG8399 | Cell membrane | 0.7929 | 0.0928 | 0.0272 | 0.0807 |
| XP_021919699.1_Zn | CG8399 | Cell membrane | 0.8736 | 0.0068 | 0.0093 | 0.1091 |
| XP_015836986.1_Tc | CG8399 | Cell membrane | 0.7084 | 0.1181 | 0.0248 | 0.1427 |
| XP_314066.4_Ag | CG8399 | Cell membrane | 0.7869 | 0.1533 | 0.0153 | 0.0371 |
| XP_026481553.1_Cf | CG8399 | Cell membrane | 0.7945 | 0.1173 | 0.0141 | 0.0683 |
| XP_021939496.1_Zn | Nemy | Lysosome | 0.2449 | 0.1184 | 0.2518 | 0.3468 |
| XP_008198104.1_Tc | Nemy | Lysosome | 0.2305 | 0.1041 | 0.2183 | 0.4366 |

| | | | | | | |
|---|---|---|---|---|---|---|
| XP_026473332.1_Cf | Nemy | Lysosome | 0.2066 | 0.1602 | 0.2203 | 0.4121 |
| XP_002430226.1_Ph | Nemy | Lysosome | 0.2073 | 0.0679 | 0.1966 | 0.52 |
| NP_001298968.1_Px | Nemy | Lysosome | 0.2463 | 0.1244 | 0.1888 | 0.4396 |
| NP_001163128.1_Dm | Nemy | Lysosome | 0.1882 | 0.1741 | 0.264 | 0.373 |
| XP_314126.2_Ag | Nemy | Lysosome | 0.1955 | 0.1699 | 0.2699 | 0.3641 |
| XP_001949276.1_Ap | Nemy | Lysosome | 0.3695 | 0.0729 | 0.0951 | 0.4533 |
| NP_001155374.1_Ap | CG1275 | Lysosome | 0.1501 | 0.0912 | 0.2134 | 0.5447 |
| XP_001950854.1_Ap | CG1275 | Lysosome | 0.1826 | 0.1045 | 0.2396 | 0.4725 |
| NP_001280323.1_Ap | CG1275 | Lysosome | 0.2415 | 0.0923 | 0.1549 | 0.5108 |
| XP_003246890.1_Ap | CG1275 | Lysosome | 0.2196 | 0.0728 | 0.2247 | 0.4798 |
| XP_008194670.1_Tc | CG1275 | Lysosome | 0.2689 | 0.0246 | 0.1127 | 0.5917 |
| XP_006572086.1_Am | CG1275 | Lysosome | 0.1834 | 0.0678 | 0.1523 | 0.5951 |
| XP_026462102.1_Cf | CG1275 | Lysosome | 0.1678 | 0.0711 | 0.1694 | 0.5907 |
| XP_021935166.1_Zn | CG1275 | Lysosome | 0.1792 | 0.1226 | 0.1842 | 0.5133 |
| XP_002426701.1_Ph | CG1275 | Lysosome | 0.3033 | 0.093 | 0.1123 | 0.4891 |
| XP_315519.4_Ag | CG1275 | Lysosome | 0.3231 | 0.0566 | 0.1535 | 0.4559 |
| NP_995963.1_Dm | CG1275 | Lysosome | 0.3271 | 0.0744 | 0.0792 | 0.5166 |

[1]The color coding was done in Excel using the Green-Yellow-Red color scale (with green for the highest value, yellow for the midpoint/50th percentile value, and red for the lowest value) across the whole data set (all four columns).
[2]ER is endoplasmic reticulum.
